# Csx15 is a broad specificity ring nuclease bridging CRISPR-Cas and CBASS cyclic nucleotide signaling systems

**DOI:** 10.64898/2026.10.26.761840

**Authors:** Danas Klimavicius, Dominik Rafalski, Dalia Smalakyte, Audroe Ruksenaite, Giedrius Sasnauskas, Giedre Tamulaitiene, Česlovas Venclovas, Matthias Bochtler, Gintautas Tamulaitis

## Abstract

Prokaryotic antiviral type III CRISPR-Cas and CBASS systems use cyclic nucleotides as second messengers to activate diverse effector proteins. Whereas CBASS signaling typically triggers abortive infection, type III CRISPR-Cas signaling is tightly regulated through controlled signal production and degradation. Here, we define the molecular mechanism of the CRISPR-associated Csx15 protein family and demonstrate that Csx15 functions as a broad specificity ring nuclease capable of degrading cyclic nucleotide signals from both CRISPR-Cas and CBASS pathways. Structural and biochemical analyses reveal that Csx15 adopts diverse oligomeric states and undergoes ligand-induced conformational transitions that promote filament assembly required for catalytic activity, analogous to filament-forming SAVED-domain effectors. Finally, we provide evidence that Csx15 is a close relative of SAVED domains suggesting that Csx15-like proteins may have also contributed to the evolution of CBASS cyclic nucleotide signaling.

## INTRODUCTION

Prokaryotes have evolved a diverse arsenal of defense systems to protect themselves from viral infection. Among these, type III CRISPR-Cas (clustered regularly interspaced short palindromic repeats; CRISPR-associated) and CBASS (cyclic nucleotide-based antiphage signaling system) employ a remarkably similar signaling strategy. In both systems, detection of phage infection triggers the synthesis of cyclic nucleotide second messengers, which subsequently activate downstream effector proteins that execute the antiviral response.

CBASS operons typically encode 2–4 proteins, including a cGAS/DncV-like nucleotidyltransferase that detects phage replication and synthesizes cyclic di-or trinucleotide signals^1^. These molecules are recognized by CBASS-associated effector proteins (Caps), which consist of a nucleotide-sensing domain fused to an effector domain. The most common sensor domains are SAVED (second messenger oligonucleotide or dinucleotide synthase-associated and fused to various effector domains) and STING (stimulator of interferon genes)^2^. However, several CBASS effectors display atypical architectures, including Cap15, which contains unique sensor domains^3^, and NucC, which lacks a dedicated nucleotide-binding domain^4^. Upon activation by cyclic nucleotide binding, Cap effectors induce abortive infection through diverse mechanisms, including nucleic acid degradation, depletion of essential metabolites, and disruption of the cell membrane^1,2^. Although additional accessory proteins (e.g., Cap2 and Cap3) may modulate synthase activity^5,6^, CBASS activation generally leads to rapid cell death, thereby restricting viral propagation within the population^2^.

Type III CRISPR-Cas systems employ a fundamentally different sensing mechanism but generate a similar signaling output^7^. These systems assemble multi-subunit ribonucleoprotein complexes composed of Cas proteins and CRISPR RNAs (crRNAs). Recognition of complementary target RNA activates the Cas10 subunit, which exhibits two catalytic activities: (i) DNase activity that degrades transcriptionally active viral DNA^8,9^, and (ii) cyclase activity that produces cyclic oligoadenylates (cA_n_; n=3-6) or related signaling molecules, such as SAM-AMP^10–12^. These second messengers accumulate and activate diverse auxiliary effector proteins that contribute to antiviral defense^13^. Most CRISPR-associated effectors contain a CARF (CRISPR-associated Rossmann fold) domain responsible for signal recognition, although SAVED-domain proteins are also found in association with type III systems.

Despite different sets of proteins, CBASS and type III CRISPR-Cas systems share a common class of signaling proteins: SAVED-domain effectors. SAVED domains were first identified through comparative genomic analyses^14^ and are currently thought to have evolved from CARF proteins through gene duplication and fusion events^15^. However, SAVED and CARF proteins differ substantially in domain architecture, ligand recognition, oligomeric state, and active site organization. Specifically, SAVED proteins typically recognize asymmetric cyclic nucleotide ligands, frequently assemble into higher-order filaments, and exhibit asymmetric active sites^15–18^. In contrast, CARF proteins typically bind symmetric ligands and function as symmetric dimers^19,20^. These structural and mechanistic differences suggest that the evolutionary origin of SAVED domains may differ from the current model.

Early models proposed that activation of type III CRISPR-Cas auxiliary effectors results in dormancy or programmed cell death, analogous to CBASS-mediated abortive infection^10^. However, signal production by Cas10 is tightly regulated and coupled to target RNA binding, with rapid termination following RNA cleavage^10,21^. Accumulating evidence indicates that type III systems primarily promote reversible growth arrest rather than irreversible cell death, thereby enhancing viral clearance while preserving host viability^22,23^. Consistently, specialized enzymes that degrade cA_n_ signals have been identified, providing a mechanism to terminate effector activation and restore cellular homeostasis^24^.

These enzymes, termed as CRISPR ring nucleases (Crn), constitute a critical regulatory layer in type III immunity. Five major classes, Crn1-Crn5, have been described. Most Crn proteins function as dimers^7^, whereas Crn3 assembles into an active complex composed of two “head-to-tail” dimers^25^. Crn1 contains a CARF domain with two conserved motifs: motif I mediates catalysis, while motif II is involved in ligand binding^24^. In contrast, Crn2-Crn5 are structurally unrelated to CARF and employ distinct architectures for cA_n_ recognition^25–28^. Substrate specificity varies among families: Crn1-3 and Crn5 preferentially degrade cA_4_^24–26,28^, whereas Crn4 exhibits broad specificity towards cA_n_^27^.

In some auxiliary effectors, including Csm6 and Cami1, the CARF domain integrates sensing and catalytic functions, degrading cA_n_ molecules to provide intrinsic negative feedback. Csm6 degrades cA_4_ or cA_6_^29,30^, whereas Cami1 preferentially targets cA_4_^23^. Similarly, the SAVED-domain-containing Lon protease CalpL functions as a ring nuclease that specifically degrades cA_4_^18,31^. Notably, ligand binding induces SAVED-dependent filament formation of CalpL, which is required for catalytic activity, revealing an additional layer of structural regulation^18^.

Bioinformatic analyses have identified a CRISPR-Cas-associated Csx15 protein family with limited sequence similarity to both CARF and SAVED domains, suggesting a potential role as ring nucleases^32^. However, experimental characterization of a Csx15 homolog from *Pseudomonas fluorescens* (PfCsx15) did not detect activity against tested cyclic oligoadenylates^33^, leaving its function unresolved. Here, we investigate the molecular mechanism and evolutionary origin of Csx15 proteins associated with type III CRISPR-Cas systems. We demonstrate that Csx15 functions as an efficient ring nuclease with broad substrate specificity, capable of degrading cyclic nucleotide signals produced by both CRISPR-Cas and CBASS pathways. We further show that Csx15 proteins adopt diverse oligomeric states and undergo ligand-induced structural transitions that promote filament formation. Structural analyses using X-ray crystallography and cryo-electron microscopy reveal that Csx15 forms filamentous assemblies required for catalytic activity, analogous to SAVED-domain effectors. Finally, we establish an evolutionary relationship between Csx15 proteins and the SAVED domain, providing evidence that SAVED proteins associated with both CRISPR-Cas and CBASS systems likely evolved from Csx15-like ancestors rather than directly from CARF proteins.

## RESULTS

### The Csx15 family consists of a major clade and several minor divergent groups

To explore the abundance and diversity of the Csx15 protein family, we initially performed sequence searches followed by sequence similarity-based grouping of identified homologs. Csx15 homologs were retrieved from the clustered non-redundant NCBI database using PSI-BLAST search with the protein from *Caldanaerobacter subterraneus* subsp. *tengcongensis* MB4 (CsCsx15) (NCBI: WP_011026657^32^) as a query. After removing sequence fragments, the remaining 764 Csx15 homologs were grouped by sequence similarity using CLANS. This analysis revealed that the majority of Csx15 homologs form a single large cluster (93% of sequences) accompanied by four minor ones (Figure 1A; Tables S1 and S2). We found that the major cluster is dominated by homologs having only the Csx15-specific region, with several additional small groups (<20 sequences each) featuring fusions N-terminal to the Csx15 region. These fusions include Schlafen, Csx1, EAD, HTH domains and transmembrane helices (TMHs) (Figure 1B).

**Figure 1.**
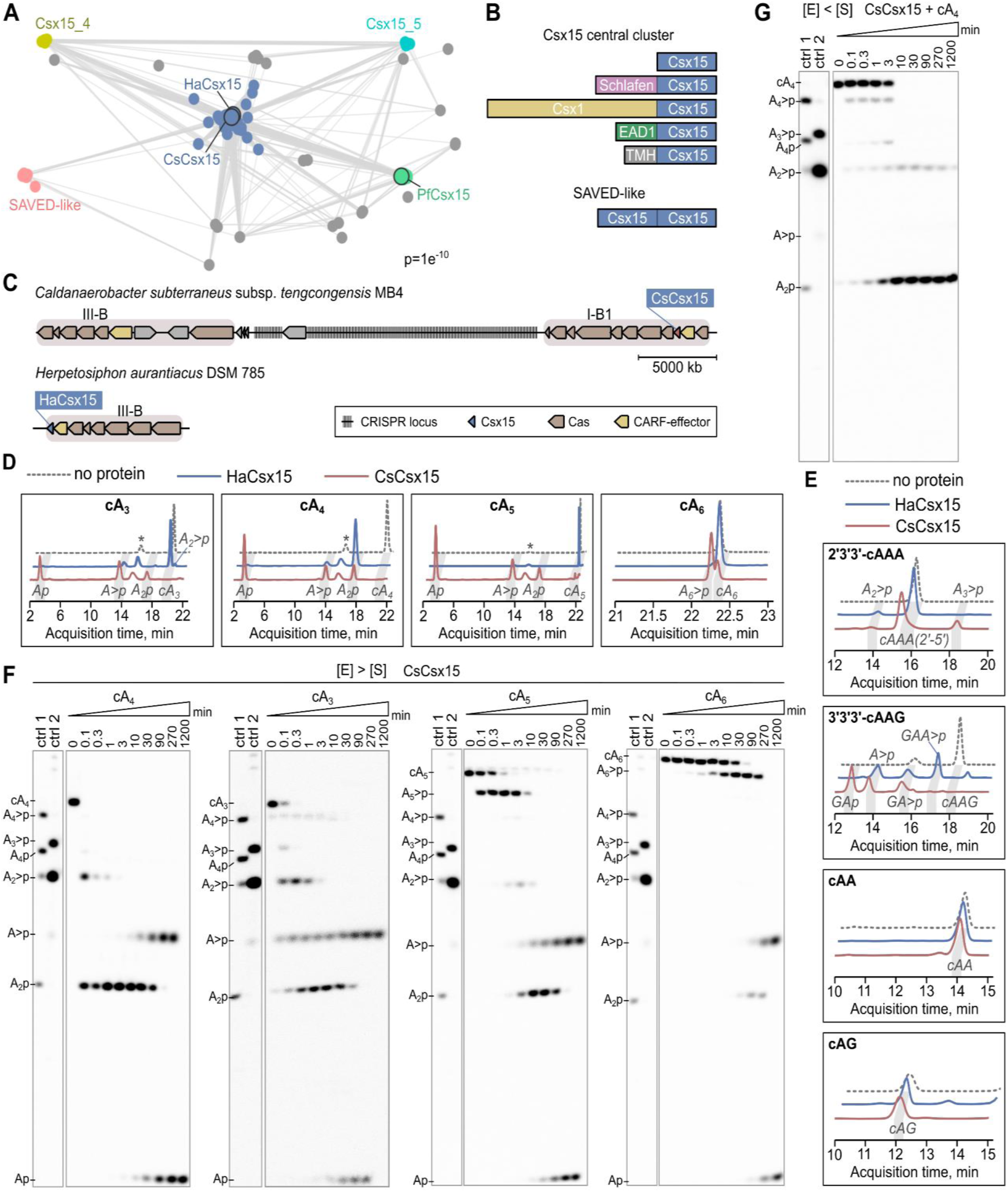
Ring nuclease activity of Csx15 on CRISPR-Cas and CBASS signaling molecules. (A) CLANS analysis of the Csx15 family. (B) Domain organization of Csx15 major cluster, shown in (A). (C) Schematic representation of genomic neighborhoods of Csx15 selected for characterization. (D, E) Hydrolysis products of CRISPR-Cas (D) or CBASS (E) signaling molecules by WT HaCsx15 and CsCsx15, analyzed by HPLC-MS. Reactions contained 10 µM of non-labeled cA_n_ and 13 µM of HaCsx15 or CsCsx15. Reaction times were 1 min for HaCsx15-cA_4_ and CsCsx15-cA_3_/cA_4_/cA_5_/3′3′3′-cAAG, 1 h for HaCsx15-cA_3_/cA_5_/cA_6_ and CsCsx15-cA_6_/2′3′3′-cAAA, 2 h for both enzymes with 3′3′-cAA or 3′3′-cAG, and 4 h for HaCsx15 with 2′3′3′-cAAA or 3′3′3′-cAAG. Asterisk (*) denotes a buffer-derived peak. (F) Cleavage of ^32^P-labeled cA_4_, cA_3_, cA_5_ and cA_6_ by WT CsCsx15 under single turnover conditions. (G) Cleavage of ^32^P-labeled cA_4_ by WT CsCsx15 under multiple turnover conditions. Reactions containing 500 nM ^32^P-labeled cA_n_ and 5 µM (single-turnover) or 50 nM (multiple-turnover) of protein were analyzed by PAGE. cA_4_ cleavage by CCaCalpL (ctrl1) and cA_6_ cleavage by StCsm6 (ctrl2) served as markers for identification of Csx15 reaction intermediates and products.

Among the four small clusters, three represent *bona fide* Csx15 proteins, including the one comprising sequences closely related to PfCsx15, which was shown to be inactive as a ring nuclease^33^. Interestingly, the remaining small cluster corresponds to proteins with a duplicated Csx15 domain (Figure S1). DALI search with an AlphaFold3 model of a representative from this group (NCBI: OGC55551) against PDB identified viral SAVED (vSAVED) ring nuclease AcrIII-2^26^ as the closest structural match (DALI Z-score: 12.6) (Figure S2A) suggesting that the SAVED domain might have evolved from the Csx15 duplication.

To investigate Csx15 structure and function, we selected two representatives featuring only the Csx15 region from the most abundant central cluster. Specifically, we selected CsCsx15 and a putative Csx15 protein from *Herpetosiphon aurantiacus* DSM 785 (HaCsx15) (Figure 1C).

The complete genome of *C. subterraneus* encodes Type III-B and Type I-B CRISPR-Cas systems, each accompanied by a CARF-domain protein gene lacking motifs characteristic of ring nucleases. Interestingly, the CsCsx15 gene is located in the vicinity of a Type I-B CRISPR-Cas system (Figures 1C and S2B), suggesting that it may regulate cA_n_ signaling *in trans*.

*H. aurantiacus* DSM 785 contains several Type III and Type I CRISPR-Cas systems (Figures 1C and S2C) and associated CARF proteins. HaCsx15 is encoded adjacent to a Type III-B system and next to a Can2/Card1 homolog, CARF domain of which lacks the motif I typical for CARF ring nucleases^34,35^. Based on this genomic context, we hypothesized that HaCsx15 may regulate Type III cA_n_ signaling *in cis* or *in trans*. CsCsx15 and HaCsx15 were cloned, expressed, and purified to homogeneity (Figure S2D). We then investigated their substrate specificity and the molecular mechanisms by which Csx15 proteins could modulate type III CRISPR-mediated antiviral immunity.

### Csx15 displays broad ring nuclease activity toward CRISPR-Cas and CBASS signaling molecules

Substrate specificity of HaCsx15 and CsCsx15 was assessed using cA_n_ of varying ring sizes (cA_3_, cA_4_, cA_5_, and cA_6_). High-performance liquid chromatography coupled to mass spectrometry (HPLC-MS) analysis showed that HaCsx15 enzyme effectively cleaves cA_4_. The major final product was A_2_p (ApA with a 3′-phosphate), with a minor product A>p (2′,3′-cyclic AMP) (Figure 1D). HaCsx15 showed limited activity toward cA_3_, generating A_2_>p and A>p, but did not cleave cA_5_ or cA_6_. In contrast, CsCsx15 displayed a broad substrate specificity, cleaving all tested substrates (cA_3_, cA_4_, cA_5_ and cA_6_) and producing A>p and Ap (3′-AMP) as final products (Figure 1D). Hydrolysis by both enzymes occurred with similar efficiency in the presence of Mg^2+^, Mn^2+^, or EDTA (Figure S3A), indicating that they function as metal-independent ring nucleases.

Given this broad specificity toward cA_n_, we further tested whether HaCsx15 and CsCsx15 can degrade signaling molecules of the CBASS pathway, which produce diverse cyclic di-and tri-nucleotides^36^. Unexpectedly, both enzymes exhibited activity toward CBASS trinucleotides (Figure 1E). CsCsx15 rapidly hydrolyzed 3′3′3′-cAAG, producing A>p and mixed dinucleotides containing adenine and guanine within one minute. It also partially converted 2′3′3′-cAAA into linear A_3_>p within one hour. HaCsx15 showed slower activity, cleaving 3′3′3′-cAAG within four hours. In contrast, neither enzyme cleaved 3′3′-c-di-AMP (3′3′-cAA) or 3′3′-cGAMP (3′3′-cAG), indicating that Csx15 targets CBASS trinucleotides containing 3′-5′ phosphodiester bonds but not dinucleotides.

To further investigate the catalytic mechanism, we performed time-course experiments under both single-turnover and multiple-turnover conditions using ^32^P-radiolabeled cA_3_-cA_6_ substrates. Reaction products were resolved by denaturing PAGE and identified by comparison with cleavage patterns generated by CCaCalpL^18^ and StCsm6^29^.

Under single-turnover conditions ([E] > [S]; 5 μM Csx15 and 500 nM substrate), both enzymes converted cA_3_ and cA_4_ into final products. Notably, CsCsx15 cleaved both substrates extremely rapidly, completing the reactions in less than a minute. HaCsx15 processed cA_4_ via a linear A_4_p intermediate, which was further cleaved into A_2_p (Figure S3B). CsCsx15 generated A_4_p intermediate, which was converted into A_2_>p and A_2_p, and finally into A>p and Ap (Figure 1F). Notably, both enzymes remained highly active even at nanomolar substrate concentrations (Figures S3C and S3D). CsCsx15 also cleaved cA_5_ and cA_6_, forming A_5_>p and A_6_>p intermediates, respectively, that were subsequently processed into final products A>p and Ap (Figure 1F).

Under multiple-turnover conditions ([S] > [E]; 500 nM substrate and 50 nM Csx15), degradation of cA_n_ by both HaCsx15 and CsCsx15 was substantially slower (calculated k_catCsCsx15(cA4)_=2.44±0.45 min^-1^ *vs* k_catHaCsx15(cA4)_= 0.27±0.03 min^-1^). HaCsx15 primarily converted cA_4_ directly to A_2_p, with minimal accumulation of the A_4_p intermediate (Figure S3E). CsCsx15 processing of cA_4_ mainly yielded A_2_>p and A_2_p, with small amounts of A_4_>p and A_4_p intermediates (Figure 1G). cA_3_ was converted directly into final products without detectable linear intermediates. However, when CsCsx15 processed cA_5_ and cA_6_, the reactions stalled at the A_5_>p and A_6_>p intermediates, respectively. (Figures S3E and S3F).

Together, these results show that (i) HaCsx15 and CsCsx15 possess broad ring nuclease activity against CRISPR-Cas and CBASS cyclic oligonucleotides, (ii) cA_4_ hydrolysis proceeds through tetra- and dinucleotide intermediates, resembling the cleavage mechanism of SAVED domain of CalpL^18^, except for CsCsx15, which converts dinucleotides further into A>p and Ap, (iii) opening of the cA_n_ ring is a rapid catalytic step, (iv) initially linearized intermediates are retained by the enzyme and further cleaved prior to products release, and (v) overall enzymatic turnover is limited by a slower step occurring downstream of ring cleavage. These findings indicate that Csx15 enzymes actively degrade, rather than merely sequester cyclic oligoadenylates, contrasting with the previously proposed “sponge” model^33^.

### CsCsx15 and HaCsx15 adopt distinct oligomeric states in solution

To investigate the molecular mechanism of Csx15 proteins, we analyzed the oligomeric states of CsCsx15 and HaCsx15 using size-exclusion chromatography coupled with multi-angle light scattering (SEC-MALS) and mass photometry. Previously characterized CRISPR ring nucleases typically function as dimers. Consistent with this, SEC-MALS analysis showed that CsCsx15 mainly forms a dimer in solution in the absence of cyclic nucleotides (Figure 2A). Addition of F_4_cA_4_, a non-hydrolysable cA_4_ analog, resulted in the appearance of higher molecular weight (M_w_) species, indicating ligand-induced oligomerization (Figure 2A). Mass photometry confirmed that apo-CsCsx15 predominantly exists as a dimer, with a minor population of tetramers (Figure S4A).

**Figure 2.**
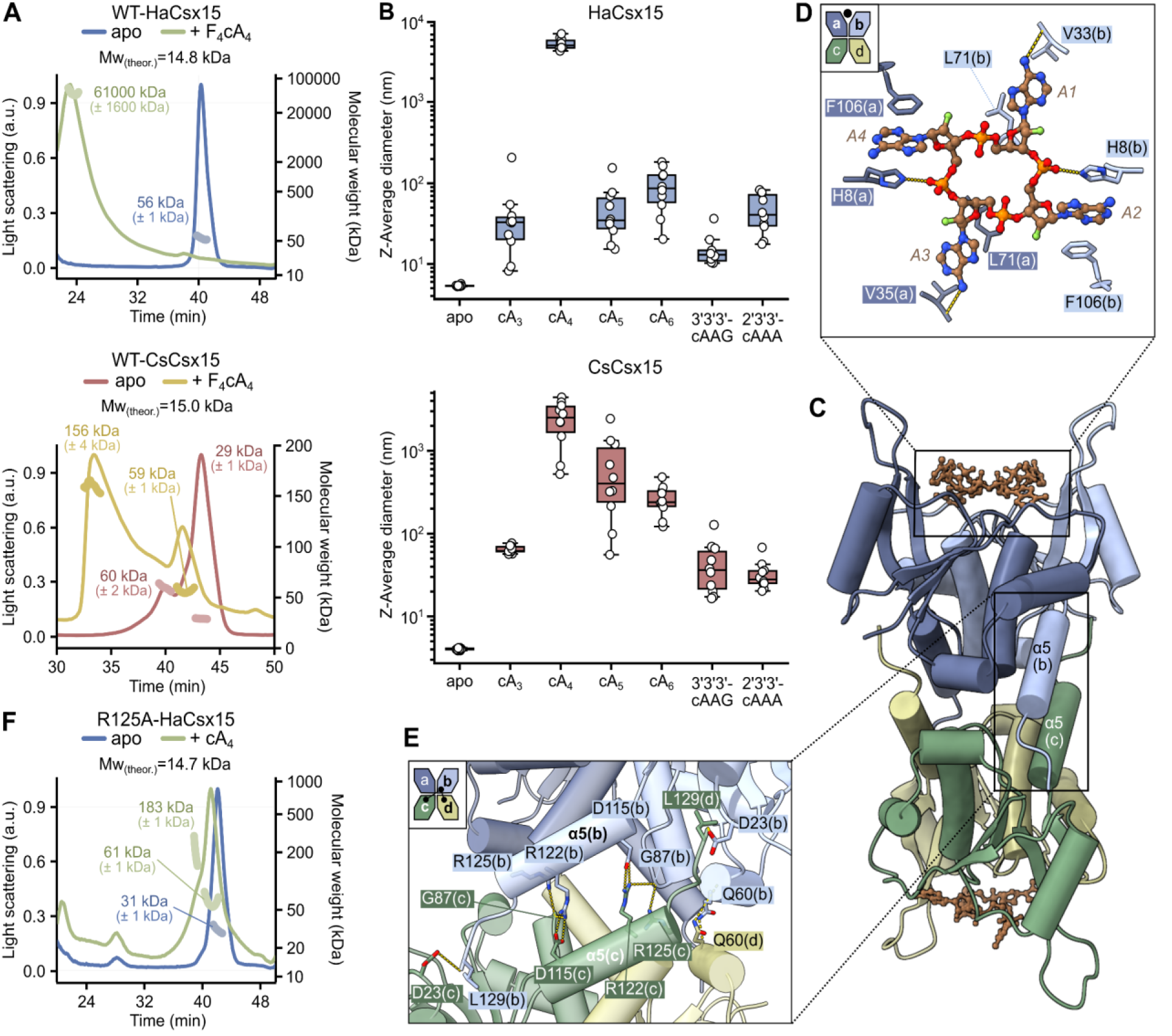
Different oligomeric states of Csx15. (A) SEC-MALS analysis of WT HaCsx15 (top) and WT CsCsx15 (bottom) in the absence or presence of F_4_cA_4_. The measured molecular weights indicate that apo HaCsx15 forms a tetramer, whereas apo CsCsx15 forms a dimer. F_4_cA_4_ binding induces assembly of higher-order oligomers in both proteins. (B) DLS analysis of cyclic nucleotide-induced oligomerization of WT HaCsx15 (top) and CsCsx15 (bottom). Data are shown as box plots (n = 10 measurements). (C) Crystal structure of WT HaCsx15 bound to F_4_cA_4_ (brown). Individual subunits are colored yellow, green, dark blue, and light blue. (D) Close-up view of HaCsx15 residues contacting the bases of F_4_cA_4_. (E) Protein-protein interactions at the HaCsx15 dimer-dimer interface. (F) SEC-MALS analysis of R125A HaCsx15, showing a molecular weight consistent with a dimeric assembly.

In contrast, HaCsx15 exhibited distinct oligomeric behavior. SEC-MALS analysis revealed that apo-HaCsx15 primarily exists as a tetramer in solution (Figure 2A). Similar to CsCsx15, addition of F_4_cA_4_, induced the formation of higher M_w_ species, consistent with ligand-dependent assembly. Mass photometry further showed that apo-HaCsx15 exists in a dimer-tetramer equilibrium (Figure S4A). Upon addition of cA_4_, larger oligomeric species appeared alongside the dimeric and tetrameric forms, and their abundance increased in a concentration-dependent manner (Figure S4B).

Bioinformatic analysis revealed that Csx15 proteins share similarity with bacterial SAVED-domain proteins^32^ and vSAVED (Figure S2A). Many SAVED-domain effectors, including the type III CRISPR-Cas effector Lon-SAVED^18^, SAVED-CHAT^37^, and CBASS effectors such as TIR-SAVED^16^, restriction endonuclease-SAVED (Cap4)^15^ and Saf-2TM-SAVED (Cap14)^17^ assemble into filamentous oligomers upon binding their activating ligands. The appearance of higher-order Csx15 species in SEC-MALS suggested that Csx15 proteins might form filamentous assemblies upon substrate binding.

To test this, we analyzed ligand-induced oligomerization of HaCsx15 and CsCsx15 using dynamic light scattering (DLS). Addition of cA_4_ to both proteins led to the formation of large particles, reaching micrometer-scale sizes (Figure 2B), indicating extensive multimerization upon prolonged exposure. As cA_4_ cleavage assays showed that CsCsx15, but not HaCsx15, converts A_2_>p and A_2_p into A>p and Ap (Figure S4C), we assumed that CsCsx15 may rebind linear dinucleotides and form oligomers. Consistently, addition of A_2_>p and A_2_p induced multimerization of CsCsx15, but not HaCsx15 (Figure S4D), confirming that CsCsx15 assembles upon binding these reaction products.

We next examined whether other CRISPR- and CBASS-derived cyclic nucleotides that serve as substrates for Csx15 ring-nuclease activity also promote oligomerization. Indeed, addition of cA_3_, cA_5_, cA_6_, and the CBASS cyclic nucleotides 3′3′3′-cAAG and 2′3′3′-cAAA resulted in the increase of particle sizes (Figure 2B). For CsCsx15, the largest assemblies were observed with cA_5_ and cA_6_, comparable to cA_4_-induced structures, whereas smaller particles formed with cA_3_, and asymmetric cyclic tri-nucleotides. HaCsx15 formed assemblies with all ligands, but these were substantially smaller than those induced by cA₄. Together, these results indicate that tetrameric HaCsx15 and dimeric CsCsx15 undergo ligand-induced assembly into higher-order oligomeric structures, potentially analogous to the filamentous assemblies observed in SAVED-domain effectors.

### HaCsx15 tetramer crystal structure

To gain structural insight into substrate recognition by Csx15 proteins, we co-crystallized HaCsx15 with F_4_cA_4_ and determined the structure of the HaCsx15-F_4_cA_4_ complex at 1.49 Å resolution (Table 1). The HaCsx15 monomer adopts a typical Rossmann-like fold, with a core composed of a five-stranded β-sheet (β1–β5) flanked by five α-helices (α1–α5) (Figure S5A).

**Table 1.** Crystallographic data collection and refinement statistics.

|  |  |
| --- | --- |
| **Structure** | **HaC<sub>s</sub>x15-F<sub>4</sub>cA<sub>4</sub>** |
| **PDB** | **30JQ** |
| **Data collection** |  |
| Resolution range(Å) | 36.37 - 1.49 (1.54 - 1.49) |
| Space group | P 3<sub>1</sub> 2 1 |
| Unit cell | 98 98 71 90 90 120 |
| Total / unique reflections | 1281074 (128937) / 58761 (4768) |
| Completeness (%) | 91.4 (75.3) |
| Mean I/sigma I | 25.5 (1.5) |
| Wilson B-factor (Å<sup>2</sup>) | 19.3 |
| R-merge (%) | 7.4 (261.9) |
| R-meas (%) | 7.6 (268.6) |
| R-pim (%) | 1.7 (59) |
| CC1/2 | 1.0 (0.78) |
| **Refinement** |  |
| R-work / R-free (%) | 15.6 (30.8) / 18.9 (35.6) |
| Number of non-hydrogen atoms (macromolecules / ligands / solvent) | 2731 (2289/87/355) |
| Protein residues | 260 |
| RMS (bonds / angles) (Å / °) | 0.007 / 1.10 |
| Ramachandran: favoured / allowed / outliers (%) | 98.44 / 1.56 / 0.00 |
| Rotamer outliers (%) | 3.72 |
| Average B-factor (Å<sup>2</sup>) | 30.56 |

Data collection statistics for the highest-resolution shell are shown in parentheses.

Structural similarity searches using the DALI server confirmed homology between HaCsx15 and both SAVED and CARF domain proteins (Figure S5B).

The structure analysis revealed that HaCsx15 assembles into a tetramer composed of two dimers, with each dimer binding a single F_4_cA_4_ molecule (Figure 2C). This oligomeric state agrees with our SEC-MALS analysis and differs from previously characterized CRISPR ring nucleases, which typically function as dimers. The tetramer is further stabilized by tartrate molecules present in the crystallization buffer. Unexpectedly, the two dimers associate in a “tail-to-tail” arrangement, positioning the two F_4_cA_4_ molecules in clefts on opposite sides of the assembly. This organization differs from the “head-to-tail” architecture previously observed in the crystal structure for the vSAVED protein^26^. Instead, the overall assembly resembles homotetrameric type IIF restriction endonucleases, which form “dimers of dimers” capable of simultaneously engaging two DNA substrates^38^.

Each dimer forms a shallow binding pocket that accommodates a single F_4_cA_4_ molecule, which sits symmetrically between the subunits. Loops spanning residues 99-104, located between β4 and β5, extend outward from each monomer and do not interfere with ligand binding. Within the pocket, F_4_cA_4_ lies along the dimer surface, with the phosphate groups oriented toward the interior and the adenine bases extending into preformed pockets. Adenine bases A2 and A4 are stabilized by stacking interactions with the side chains of H8 and F106, whereas A1 and A3 are located in the pocket formed by R37 and L71 side chains and interact with the protein backbone of the residue V35 (Figure 2D).

Tetramer formation is stabilized by the cross-dimer contacts, generating a total interface area of approximately 2160 Å². Within this interface, the C-terminal helix α5 mediates key electrostatic interactions between the dimers, including contacts between R125 and R122 from one dimer and G87 and D115 residues from other dimer, respectively, as well as additional interactions between the main chain atoms of D23 and L129 (Figure 2E). To test the importance of this interface, we generated the R125A variant. We further assessed the apo oligomeric state of R125A HaCsx15 using SEC-MALS. In contrast to wild type (WT) protein, which forms tetramer, R125A HaCsx15 variant was detected predominantly as a dimer (Figure 2F). This result indicates that R125 plays a critical role in stabilizing dimer-dimer interactions.

### cA_4_-activated Csx15 assemble into filaments

To elucidate the structural basis of substrate induced multimerization of Csx15, we performed cryogenic electron microscopy (cryo-EM) analysis of HaCsx15 and CsCsx15 in the presence of different substrates (Table 2 and Figure S6). We determined the structure of HaCsx15 bound to F_4_cA_4_. For CsCsx15, we obtained two complexes, one with cA_4_ and another with 3′3′3′-cAAG, but only the former was resolved at high resolution.

**Table 2.** Cryo-EM data collection, refinement and validation statistics.

|  |  |  |
| --- | --- | --- |
| PDB<br>EMDB | **HaCsx15-F4cA4**<br>28WJ<br>EMD-56908 | **CsCsx15-A2>p**<br>30FQ<br>EMD-57705 |
| **Data collection and processing** |  |  |
| Microscope | Glacios (TFS) | Glacios (TFS) |
| Detector | Falcon III camera (TFS) | Falcon III camera (TFS) |
| Magnification | 92,000 | 92,000 |
| Automation software | EPU | EPU |
| Voltage (kV) | 200 | 200 |
| Electron exposure (e/Å<sup>2</sup>) | 29.7 | 29.7 |
| Defocus range (μm) | 0.5 to 2.5 | 0.5 to 2.5 |
| Pixel size (Å) | 1.1 | 1.1 |
| Micrographs collected/used | 835/664 | 1,178/976 |
| Initial/final particle images (no.) | 400,533/46,968 | 1,9256,725/260,938 |
| Symmetry imposed | C1 | C2 |
| Helical rise (Å)/twist (°) | 15.75/173.62 | 31.95/16.77 |
| Map resolution (Å) (masked/unmasked); FSC threshold 0.143 | 3.0/4.0 | 3.06/3.8 |
| Map sharpening B factor (Å<sup>2</sup>) | 104.4 | 179.51 |
| **Refinement** |  |  |
| Initial model used | AlphaFold3 | AlphaFold3 |
| Model resolution (Å) FSC threshold 0.5 | 3.1 | 3.2 |
| Model composition<br>Non-hydrogen atoms/Protein residues/Nucleotides | 16,576/2064/32 | 6,478/769/12 |
| B factors (mean, Å<sup>2</sup>)<br>Protein/Nucleotide | 46.00/37.61 | 60.87/62.99 |
| RMSDs<br>Bond lengths (Å) /Bond angles (°) | 0.04/0.818 | 0.06/0.778 |
| **Validation** |  |  |
| MolProbity score | 1.46 | 1.40 |
| Clash score | 5.87 | 4.72 |
| Poor rotamers (%) | 0.70 | 1.59 |
| Ramachandran plot<br>Favored/Allowed/Disallowed (%) | 97.24/2.76/0.00 | 98.15/1.85/0.00 |
| CCvolume/CCmask | 0.85/0.88 | 0.78/0.81 |
| Cb outliers (%) | 0.00 | 0.00 |
| CaBLAM outliers (%) | 1.35 | 0.94 |

The HaCsx15-F_4_cA_4_ structure revealed that HaCsx15 dimers stack into long, well-ordered double-filamentous assemblies, approximately 100 nm in length (with 32 protein dimers in each filament) (Figures 3A). From the 3.0 Å reconstruction, we modeled four pairs of Csx15 dimers sandwiching F_4_cA_4_ molecules. Dimers oligomerize in a “head-to-tail” manner, with a single activator molecule located at the interface between adjacent dimers, forming a single-stranded filament (Figure 3B). Multiple cross-strand interactions further organize these structures into a higher order double-stranded filament with a visible twist, in which the two strands are oriented in parallel. The double-strand interfilament interface is mediated by hydrogen bonds between the residues D23-R39 and N100-G109, reinforced by additional van der Waals interactions (Figure 3C).

**Figure 3.**
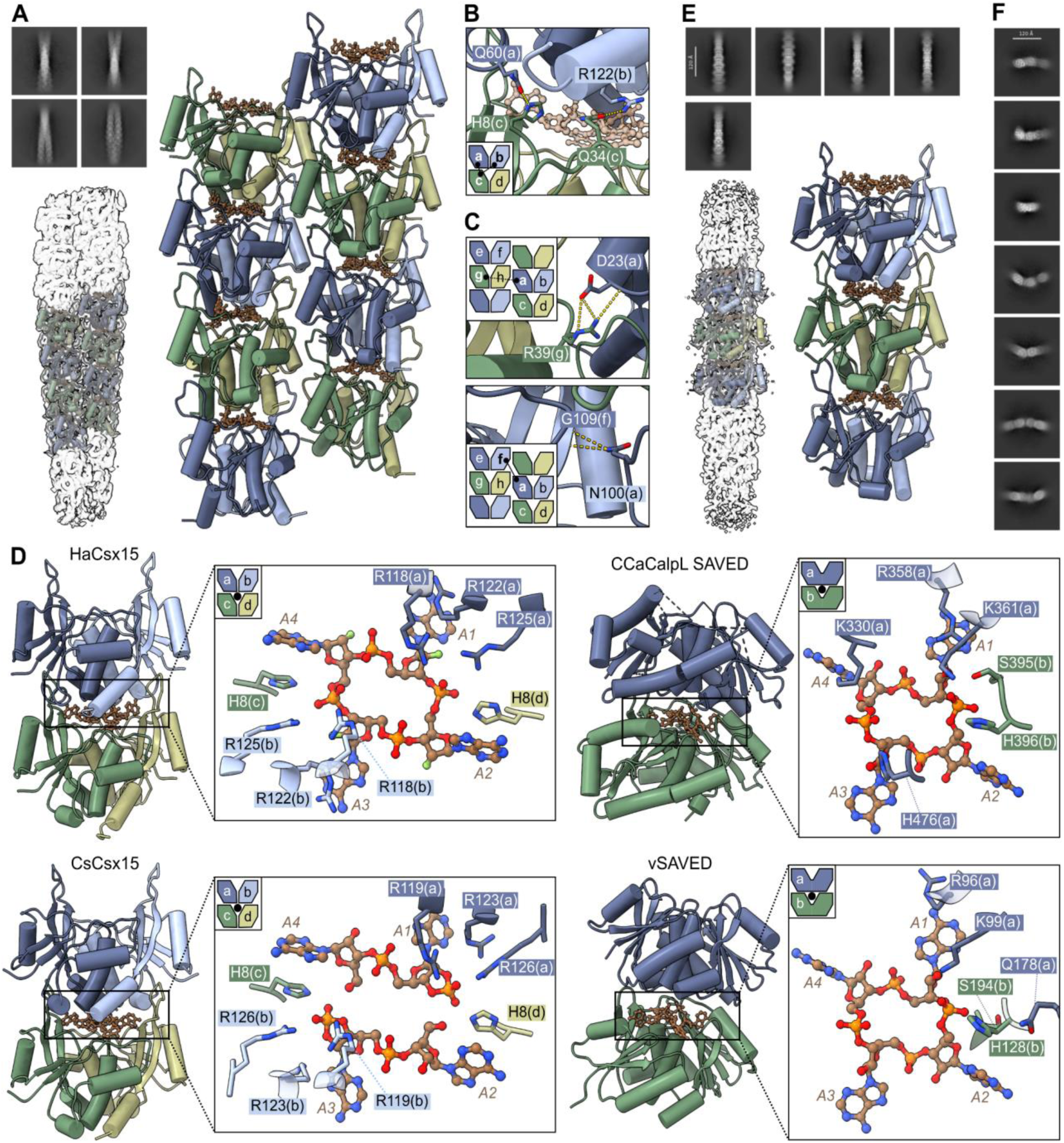
Cryo-EM structures of HaCsx15-F_4_cA_4_ and CsCsx15-A_2_>p complexes. (A, E) Representative 2D classes (top left), cryo-EM density maps (bottom left), and atomic models (right) of WT HaCsx15 bound to F_4_cA_4_ (A) and WT CsCsx15 bound to A_2_>p (E). The density maps show eight and three adjacent dimers, respectively. F_4_cA_4_ and two A_2_>p molecules, generated by cA_4_ cleavage (brown), are bound at the interfaces between neighboring dimers (colored in shades of blue and olive). (B) Amino acid interactions at the HaCsx15 dimer-dimer interface within a single filament strand. (C) Amino acid interactions mediating assembly of the double-stranded HaCsx15 filament. (D) Comparison of neighboring subunits from HaCsx15-F_4_cA_4_, CsCsx15-A_2_>p, CCaCalpL SAVED-cA_4_ (PDB 9EYJ), and vSAVED (PDB 9R8S; cA_4_ modeled). Insets show close-up views of potential catalytic residues interacting with the ligand in each complex. (F) Representative 2D class averages of WT CsCsx15 oligomers bound to 3′3′3′-cAAG.

F_4_cA_4_ is tightly sandwiched between neighboring dimers by extensive interactions contributed from both surfaces. The dimer-dimer interface itself contains relatively few direct protein-protein contacts, indicating that filament assembly is primarily stabilized through protein-substrate interactions. Contacts observed in the tetramer structure, including interactions of H8 and F106 with adenines A2 and A4, are supplemented by additional contacts from neighbouring dimers. In particular, two R122 from the “tail” surface form π-stacking interactions with adenines A1 and A3. Additional contacts to the phosphodiester backbone involve R118, R122 and R125 “tail” residues (Figure 3D). The scissile phosphates form hydrogen bonds with H8 and R125, suggesting roles in stabilization of the pentavalent transition state. Notably, H8 is positioned near the 5′-oxygen leaving group and may participate in proton transfer during catalysis.

Next we determined the cryo-EM structure of CsCsx15-cA_4_ complex at 3.06 Å resolution (Figure 3E). CsCsx15 forms a single-stranded filament composed of dimers arranged in a “head-to-tail” configuration, resembling HaCsx15 filaments in both architecture and dimensions. Within the filament, cA_4_ is cleaved to the product A_2_>p. Conserved residues H8 from the “head” surface and R119, R126 from the “tail” surface, equivalent to H8 and R118, R125 in HaCsx15, are positioned to interact with the cleavable phosphate groups (Figure 3D). Their spatial arrangement is consistent with their potential catalytic role. Differently from cA_4_, 3′3′3′-cAAG induced formation of noticeably shorter asymmetric CsCsx15 oligomers (Figure 3F).

### Ring nuclease activity relies on the Csx15 filament formation

We compared HaCsx15-F_4_cA_4_ and CsCsx15-A_2_>p filament structures with structures of the filament-forming SAVED domain proteins with ring nuclease activity - CcaCalpL bound to cA_4_ (PDB: 9EYJ) and the apo vSAVED dimer (PDB: 9R8S) (Figure 3D). In both systems, SAVED monomers adopt a “head-to-tail” arrangement and form a composite active site for cA_4_ cleavage. Conserved residues from both monomers contribute to cA_4_ binding and catalysis in a single active site formed at the dimer interface^18,26^. Both HaCsx15 and CsCsx15 dimers are structurally analogous to individual SAVED monomers of CCaCalpL and vSAVED (Figure 3D). To identify catalytic residues, we superimposed the Csx15 monomer with the C-subdomains of CcaCalpL and vSAVED. Catalytic residues H396/H128 of CCaCalpL/vSAVED correspond to HaCsx15/CsCsx15 H8 located on the “head” surface. The CcaCalpL/vSAVED active site residues R358, R361/R96, R99 lack direct equivalents in Csx15s. Instead, HaCsx15/CsCsx15 residues R118/R119, R122/R123 and R125/R126 from the “tail” surface are positioned to interact with cA_4_ phosphates.

To test the roles of these residues, we generated H8A and R118A variants of HaCsx15 and analyzed their oligomeric state and activity using SEC-MALS, DLS and ^32^P-labeled cA_4_ hydrolysis assays. Both variants remained tetrameric in the absence of cA_4_ (Figure S7A). H8A retained cA_4_-induced multimerization, indicating preserved ligand binding (Figure 4A). However, substitution of H8 with alanine abolished enzymatic activity (Figure 4B), suggesting that H8 functions as a catalytic residue. In contrast, the R118A variant lost both ring nuclease activity and the ability to multimerize, indicating that R118 is important for cA_4_ binding during filament assembly (Figure 4B).

**Figure 4.**
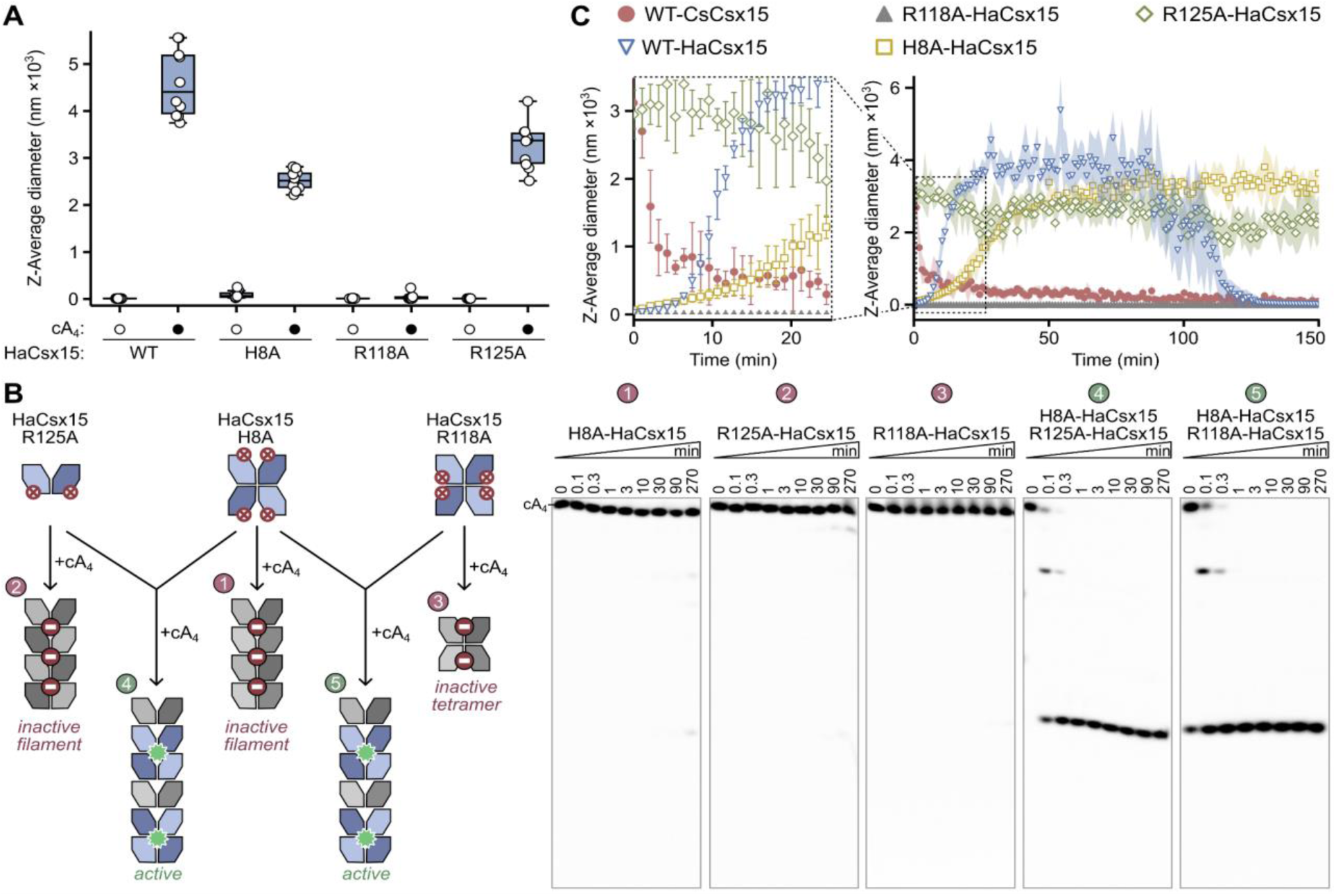
cA_4_-induced HaCsx15 rearrangement and filament assembly generate the composite active site. (A) DLS analysis of cA_4_-dependent oligomerization of WT HaCsx15 and putative active-site mutants. Data (n = 10 measurements each) are shown as box plots. (B) Schematic of composite active-site assembly in the cA_4_-activated HaCsx15 filament (left). Single-turnover ring nuclease reactions of putative active-site mutants (reactions 1-3) and filaments assembled by mixing two mutant HaCsx15 proteins (reactions 4-5), analyzed by denaturing PAGE (right). H8A and R125A abolished catalytic activity but retained cA_4_ binding and oligomerization, identifying H8 and R125 as catalytic residues. Mixing two inactive mutants restored cA_4_ cleavage to WT levels, consistent with formation of a composite active site within the filament. Reactions contained 5 µM HaCsx15 and 500 nM ^32^P-labeled cA_4_. (C) DLS kinetics of cA_4_-induced oligomerization of WT and mutant HaCsx15. Each point represents the median of n = 10 measurements. WT tetrameric HaCsx15 assembled into multimers gradually with a lag phase, whereas dimeric R125A formed multimers immediately upon cA_4_ addition.

To further examine the relationship between oligomerization and catalysis, we analyzed the R125A variant, which disrupts “tail-to-tail” tetramer formation in the apo state (Figure 2F). Although R125A forms dimers in solution, it assembles to higher-order multimers upon cA_4_ addition, as detected by DLS (Figures 4A). This indicates that filament formation does not require the “tail-to-tail” apo-tetramer interface. However, R125A lacked cA_4_ ring nuclease activity (Figure 4B), demonstrating that R125 is essential for catalysis. These results suggest a dual role for R125: stabilizing dimer-dimer contacts in the apo tetramer and contributing to formation of the composite active site in the filament together with H8 from a neighboring dimer.

To test this model, we reconstituted heterotetramers by mixing the inactive H8A and R125A variants, which carry mutations on the opposite surfaces of the HaCsx15 dimer. Mixing restored ring nuclease activity to wild-type levels (Figure 4B), consistent with the hypothesis of the formation of a composite active site within the filament. A similar recovery of activity was observed when H8A was combined with R118A, further supporting that “head-to-tail” tetramer assembly represents the minimal catalytic unit of HaCsx15 (Figure 4B).

### HaCsx15 filament formation requires tetramer rearrangement

To monitor changes in the Csx15 oligomeric state in time, we performed DLS kinetics experiments using WT and mutant variants of HaCsx15 and CsCsx15 (Figure 4C). For both WT HaCsx15 and its H8A mutant, in the presence of cA_4_ we observed gradual increase of high M_w_ multimers with their maximum sizes reaching after approximately 10-15 minutes. This delay suggests that the filament assembly requires disruption of the “tail-to-tail” tetramer into dimers, which can then reorganize into a “head-to-tail” arrangement that supports higher-order filament formation. For WT HaCsx15, but not for the ring nuclease-dead H8A mutant, the multimeric particles disappeared after ∼100 minutes of incubation. This time scale is consistent with the cA_4_ cleavage time of HaCsx15 under multiple-turnover conditions (Figure S3E) and most likely reflects filament disassembly following cA_4_ degradation. As expected, the HaCsx15 R118A mutant, defective in filament formation, did not form multimers during the measurement.

In contrast, addition of cA_4_ to the tetramer-disrupting, dimeric R125A variant resulted in immediate formation of large multimers, with no detectable lag phase. This suggests that the dimeric mutant can directly assemble into the “head-to-tail” filament architecture without first undergoing tetramer dissociation. Consistent with this interpretation, immediate multimer formation was also observed when cA_4_ was added to the naturally dimeric WT CsCsx15. Together, these results indicate that pre-existing dimers can bind cA_4_ and assemble into filaments without additional structural rearrangements, in agreement with the filament assembly kinetics observed for the dimeric HaCsx15 R125A variant.

### Filament formation reshapes the active site into an active conformation

In both the non-filamentous and filamentous forms of HaCsx15, a 2’-OH of cA_4_ would be poised for in-line nucleophilic attack on the scissile phosphate, which should cause the departure of the 5’-OH leaving group. In the crystals and the cryo-EM structure, the cleavage reaction is artificially prevented by 2’-fluoro substitution. Despite this artificial perturbation of the active site, which prevents cleavage in both cases, there is a clear difference in the active site that explains in structural terms why filament formation drives cA_4_ hydrolysis. In the non-filamentous form, the Nε is hydrogen bonded to the proR oxygen of the scissile phosphate. While this interaction helps to anchor cA_4_, it is not expected to directly facilitate catalysis. In the filament form, the histidine moves towards the leaving group, and could donate a hydrogen bond to the departing 5’-oxygen (O5’). The interaction should lower the barrier for hydrolysis by making the O5’ a better leaving group, and it also provides a mechanism for the protonation of the O5’ to the 5’-OH in the product (Figure S7B).

### Bacterial Csx15 is a functional ring nuclease *in vivo*

To investigate the *in vivo* activity of Csx15 protein, we reconstituted HaCsx15 into a recombinant type III-A CRISPR system from *Streptococcus thermophilus* (StCsm)^39^. This system generates diverse cA_n_^29^, including cA_4_, which potently activates the cA_4_-dependent AvCami1 effector, leading to growth arrest in *E. coli*^23^.

For functional assays, *E. coli* cells expressing StCsm from a three plasmid, IPTG-inducible system (pCsm, pCRISPR, pTarget (Table S4)) were transformed with a plasmid encoding the AvCami1 effector alone or co-expressing AvCami1 and with either WT or the H8A mutant HaCsx15 gene (Figure 5A). As a negative control, cells carried an pEmpty vector. In this setup, Csx15 activity was expected to counteract the growth inhibition caused by cA_4_-activated AvCami1, which is triggered by StCsm programmed to target the pTarget transcript and thereby produce the cA_4_ activator.

**Figure 5.**
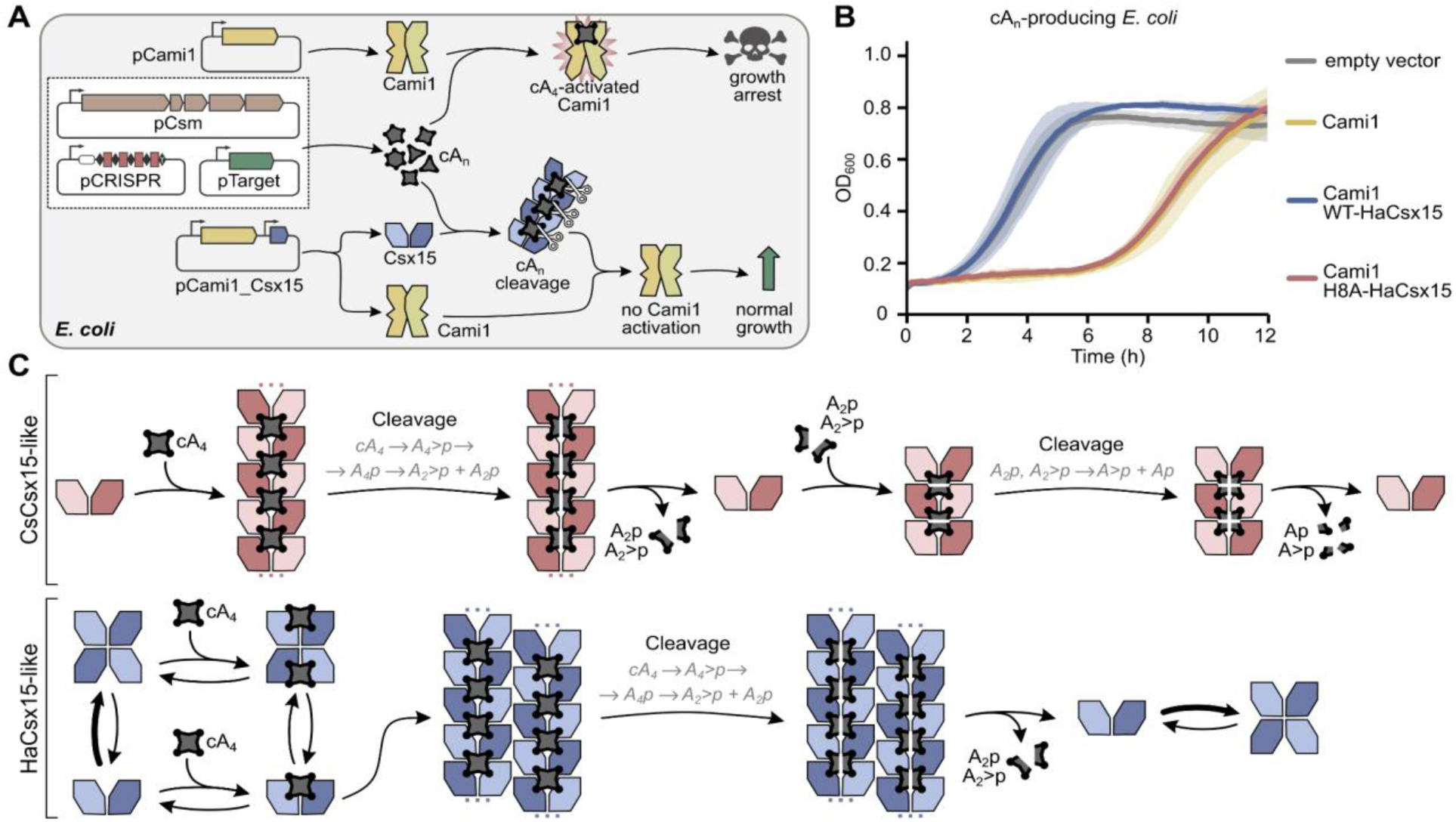
Csx15 activity *in vivo* and cA_4_ cleavage reaction mechanism. (A) Experimental design for analysis of Csx15 activity *in vivo*. (B) Csx15 reduces intracellular cA_4_ levels, as its expression in *E. coli* prevents growth arrest induced by the cA_4_-activated effector Cami1. (C) Model of cA_4_ hydrolysis by HaCsx15-like and CsCsx15-like ring nucleases. Csx15 proteins adopt distinct resting oligomeric states and undergo cA_4_-induced rearrangement to assemble into active filaments. Csx15 dimers form a composite active site within assembled filament for cA_4_ cleavage, in a manner similar to the Lon-SAVED monomer active site. Following cleavage of cA_4_ into dinucleotides, filaments disassemble and return to the resting state. Upon rebinding of the cleavage products, CsCsx15-like filaments can further process dinucleotides into mononucleotides.

In the absence of HaCsx15, *E. coli* exhibited pronounced growth defects upon the induction of AvCami1 and cA_n_ synthesis (Figure 5B). In contrast, expression of WT HaCsx15 restored culture growth to the level of the negative control, demonstrating that HaCsx15 effectively reduces intracellular cA_4_ concentrations. Mutation in the HaCsx15 active site (H8A variant) impaired this protective effect, resulting in strong growth inhibition for HaCsx15-H8A, confirming the essential role of the H8 residue in HaCsx15 catalytic activity. Together, these findings demonstrate that HaCsx15 protein is a functional ring nuclease *in vivo* and effectively reduces the intracellular cA_4_ concentration.

### Csx15 co-occurrence with CBASS in bacterial genomes

The finding that Csx15 could hydrolyze both cA_n_ and selected CBASS messengers prompted us to investigate its genomic association with CRISPR-Cas and CBASS systems. We analyzed 1,204 bacterial and metagenomic genomes from NCBI containing at least one annotated Csx15 protein and ≥2000 predicted open reading frames. Csx15 proteins were distributed across diverse bacterial phyla, with enrichment in *Chloroflexota*, *Planctomycetota*, and *Pseudomonadota* (Figure 6A and Table S3). We then screened these genomes for homologs of Cas10, the CRISPR-associated cA_n_ synthase, and the CBASS synthases CdnD (3′3′3′-cAAG, 2’3’3’-cAAA synthase)^15,40^, CdnE (3’3’-cUA, 3′3′-cUU, 3′3′-cGAMP, 3′3′-c-di-GMP synthase)^36,40^, and DncV (3′3′-cGAMP synthase)^41^.

**Figure 6.**
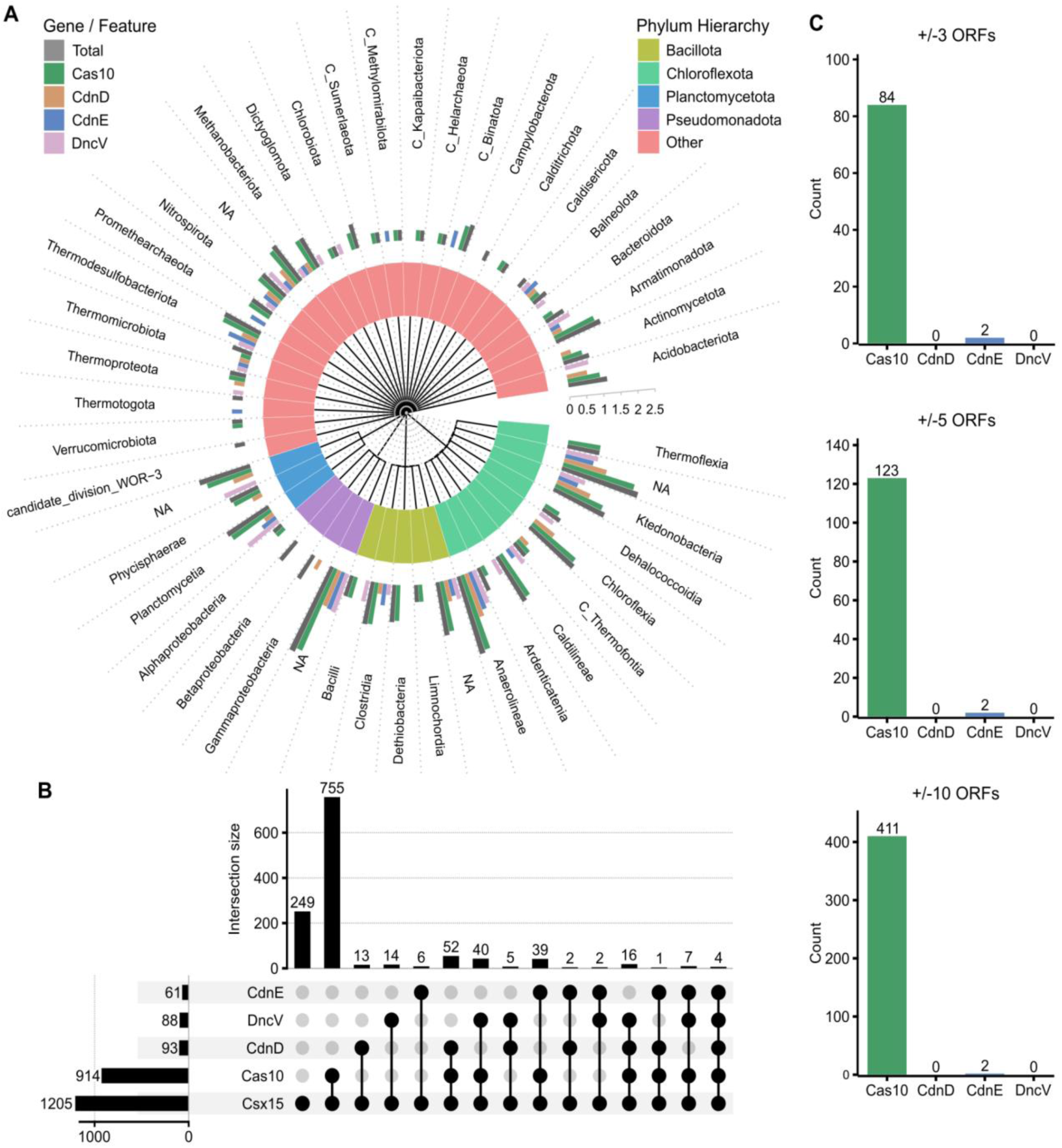
Association of Csx15 with CRISPR-Cas and CBASS signaling systems. (A) Species counts according to phylogeny. For each leaf of the phylogenetic tree, the number of species with a CRISPR-Cas synthase (Cas10) and CBASS synthases (CdnD-like, CdnE-like, DncV-like) was counted. The log_10_(<NUMBER of species>+1) is plotted. C stands for “Candidatus”. (B) Upset plot for the co-occurrence of Csx15 with Cas10-, CdnD-, CdnE-, and DncV-like proteins. Note that only metagenomes with Csx15 were considered, hence Csx15 is universally present. (C) Genomic neighborhoods of Csx15.

Across phyla, Csx15 most frequently co-occurred with Cas10 (∼80%), consistent with its established role in type III CRISPR-Cas signaling (Figure 6B and Table S3). Co-occurrence with CBASS synthases was less common (∼20%) and often accompanied by Cas10, precluding direct assignment of a CBASS-specific function. However, ∼4% of genomes encoded Csx15 together with one or two CBASS synthases but lacked Cas10, supporting a potential independent role of Csx15 in CBASS signaling. Among CBASS-associated systems, CdnD was the most prevalent synthase, notable because it produces 3′3′3′-cAAG, one of the most efficiently degraded substrates of Csx15.

Because genomic proximity can indicate functional association, we examined genes located near *csx15* loci. Cas10 was frequently found in the immediate genomic neighborhood of Csx15, further supporting coupling with type III CRISPR-Cas systems (Figure 6C and Table S3). In contrast, CdnD, CdnE, and DncV were rarely adjacent to *csx15* genes. These observations suggest that, unlike CRISPR-associated Csx15 proteins, CBASS-associated Csx15 proteins may not require stable genomic linkage to synthase genes, consistent with the *in trans* activity observed for CsCsx15.

### Csx15 homologs in viruses

During search for Csx15 homologs in the NCBI ClusteredNR database, we detected only a single viral homolog, a hypothetical protein (NCBI: URC17498) termed as vCsx15 derived from the MAG genome of Lokiarchaeota virus Skoll Meg22_1214^42^. Structural modeling with AlphaFold3 indicated that this protein indeed showed close structural similarity to the structure of prokaryotic HaCsx15 protein (Figure S8). We hypothesized that found viral vCsx15 may function as viral ring nucleases. Therefore, we tentatively propose the name AcrIII-3 (anti-CRISPR ring nuclease 3) for this family of proteins, joining characterized AcrIII-1 and AcrIII-2 families^24,28^.

To determine whether additional viral Csx15 homologs exist, we used BLAST to search the NCBI database of proteins from WGS metagenomic projects (env_nr) and the IMG/VR database of metagenomic viral protein sequences. In total we identified over 20 viral sequences (an aligned subset is shown in Figure S8). Importantly, all identified sequences originate from metagenomic datasets and the functional activity of these proteins has yet to be demonstrated experimentally.

## DISCUSSION

### Mechanism of Csx15 ring nucleases

In this study, we show that type III CRISPR-Cas accessory Csx15 proteins form a broad conserved family of ring nucleases that degrade cyclic signaling molecules both *in vitro* and *in vivo*. These enzymes can act *in cis* or *in trans* within bacterial cells suggesting that it actively participates in regulation of type III CRISPR-Cas immunity. Unlike previously characterized Crn1-Crn3 nucleases^24,25,28^, Csx15 proteins display broad substrate specificity, cleaving cA_3_-cA_6_. Importantly, Csx15 also hydrolyzes CBASS signaling molecules, including 3′3′3′-cAAG and 2′3′3′-cAAA, revealing an unexpected functional overlap between CRISPR-Cas and CBASS pathways. This broad specificity, together with structural features, further links Csx15 proteins to SAVED domains found in CBASS effectors.

Our structural and biochemical analyses reveal that Csx15 proteins employ a catalytic strategy that is conceptually similar to that of Lon-SAVED effectors. In both systems, ring nuclease activity depends on ligand-induced assembly into filamentous oligomers that generate composite active sites at subunit interfaces. However, the pathways leading to filament formation differ substantially among Csx15 proteins. As illustrated in Figure 5C, apo-CsCsx15 predominantly exists as a dimer in solution, resembling previously characterized Crn1-Crn4 ring nucleases. Binding of cyclic nucleotide substrates promotes assembly of these dimers into “head-to-tail” filaments, thereby creating the composite active sites required for catalysis. In contrast, HaCsx15 exists in a dimer-tetramer equilibrium, with the tetramer representing the dominant species. The tetramer adopts a “tail-to-tail” architecture that appears to correspond to an inactive resting state. Upon substrate binding, this tetramer likely dissociates into cA_4_-bound dimers, which subsequently reorganize into a “head-to-tail” arrangement and assemble into catalytically active filaments. Notably, residue R125 performs a dual role in this process. In the apo state, it stabilizes the inactive tetramer through dimer-dimer interactions, whereas in the filament it contributes directly to formation of the composite active site. Based on this model, the crystal structure of HaCsx15 represents an intermediate state in the activation pathway, capturing a substrate-bound tetramer poised for dissociation and subsequent rearrangement into an active filament. Filament assembly not only generates the composite active site but also remodels it into a catalytically competent conformation, positioning H8 for efficient substrate cleavage.

Although both Csx15 and Lon-SAVED enzymes rely on filament formation for activation, their catalytic mechanisms differ. Csx15 dimers contribute two symmetric active sites per subunit pair, enabling concerted cleavage of cA_4_ within the filament. In contrast, Lon-SAVED enzymes contain a single active site per monomer and process cA_4_ through a stepwise mechanism involving repeated substrate dissociation and rebinding. Under multiple-turnover conditions, Lon-SAVED catalyzes cA_4_ conversion via a sequential three-step pathway, leading to accumulation of A_4_>p and A_4_p intermediates before formation of A_2_p and A_2_>p. In contrast, Csx15 predominantly converts cA_4_ directly to A_2_>p and A_2_p, with minimal intermediate accumulation. Consistent with this, Csx15 forms extended, stable filaments that support efficient and processive catalysis, while Lon-SAVED forms shorter, transient assemblies. Furthermore, at least in the case of CsCsx15, the resulting A_2_>p and A_2_p products can be further degraded to A>p and Ap, a capability not observed for Lon-SAVED enzymes. Following substrate cleavage, filaments disassemble back into their resting states: dimers for CsCsx15 and tetramers for HaCsx15.

The activation mechanism proposed for HaCsx15, involving a tetramer-dimer-filament transition, resembles the recently described activation pathway of SPARDA^43^. In both systems, an inactive oligomeric assembly must first dissociate before productive higher-order complexes can form.

This multistep activation pathway likely serves as an additional safeguard against spontaneous or unintended ring nuclease activation. Notably, both CsCsx15 and HaCsx15 remain catalytically active at nanomolar concentrations, indicating efficient filament assembly even at low intracellular protein abundance. This supports a physiological role for Csx15 in clearing residual cA_n_ following infection and limiting unintended signaling. During type III CRISPR-Cas immune responses, the interference complex generates a spectrum of cA_n_. In this context, the broad substrate specificity of Csx15 supports its role as a versatile ring nuclease involved in regulating type III CRISPR-Cas immunity.

Recent work by Chi *et al*. reported that Csx15 from *Pseudomonas fluorescens* and *Chlorobaculum limnaeum* form “head-to-tail” dimer stacks in crystal lattices and assemble into higher-order oligomers in solution^44^. While PfCsx15 lacks detectable activity and possibly represents a non-functional outlier (Figure 1A), dimeric CliCsx15 cleaves cA_4_ at micromolar concentrations and may follow a CsCsx15-like activation mechanism. Together, these findings support a general model in which filament formation is essential for ring nuclease activity across the Csx15 family, despite variation in their resting-state oligomeric organizations and activation pathways.

### Csx15 as the likely evolutionary predecessor of SAVED domain

An evolutionary relationship between CARF and SAVED domains previously led to the hypothesis that pseudosymmetric SAVED domains arose through fusion of two CARF monomers^15,32^. However, our results prompt a revision of this scenario, identifying Csx15 as a more likely evolutionary predecessor of the SAVED domain. Previous sequence analysis suggested that Csx15 is related to both CARF and SAVED^32^. Our results reported here not only confirm this proposition, but also establish that Csx15, and not CARF, is structurally closer to SAVED (Figure 7). In particular, compared to CARF, Csx15 and each SAVED subdomain lack one of the α-β motifs. In addition to close structural similarity, Csx15 and SAVED share a number of other features not seen in CARF proteins. CARF domains possessing the ring nuclease activity have two active site motifs on the same side of the protein surface^24^. In contrast, Csx15 and those SAVED domains that are capable of degrading signaling molecules have a composite active site that involves residues located on the opposite faces of a monomer^18^. The active site becomes fully assembled only upon stacking of Csx15 dimers or SAVED monomers in a “head-to-tail” oligomeric filament with cyclic nucleotides trapped at the interfaces. The differences in the active site organization perhaps might explain why CARF ring nucleases have not been observed to form filaments similar to those of Csx15 and SAVED. Therefore, a recent report of TIR-CARF protein dimers forming stacked filaments^45^ appeared to be an exception. However, upon closer examination we found that what was identified as a CARF domain^45^, is in fact a domain of the Csx3 family exemplified by the ring nuclease Crn3 also forming “head-to-tail” oligomers^25^. Another shared feature of SAVED and Csx15 proteins is the ability to recognize multiple cyclic nucleotides, including asymmetric ones. While such promiscuity is expected for SAVED domains due to their pseudosymmetric architecture, it is an unexpected feature for symmetric Csx15 dimers. This functional flexibility further supports a model in which SAVED domains evolved via fusion of two Csx15-like units followed by sequence and structure adjustment. In support of such a scenario, we identified proteins containing Csx15-Csx15 fusions (Figure 1B). Interestingly, these fused proteins often lack residues required for one or both active sites. This might be a general feature of pseudosymmetric ring nucleases due to the relaxed evolutionary pressure compared with symmetric dimers formed by monomeric proteins. In agreement with this view, all SAVED ring nucleases characterized to date contain a single functional active site^18,26,31^.

**Figure 7.**
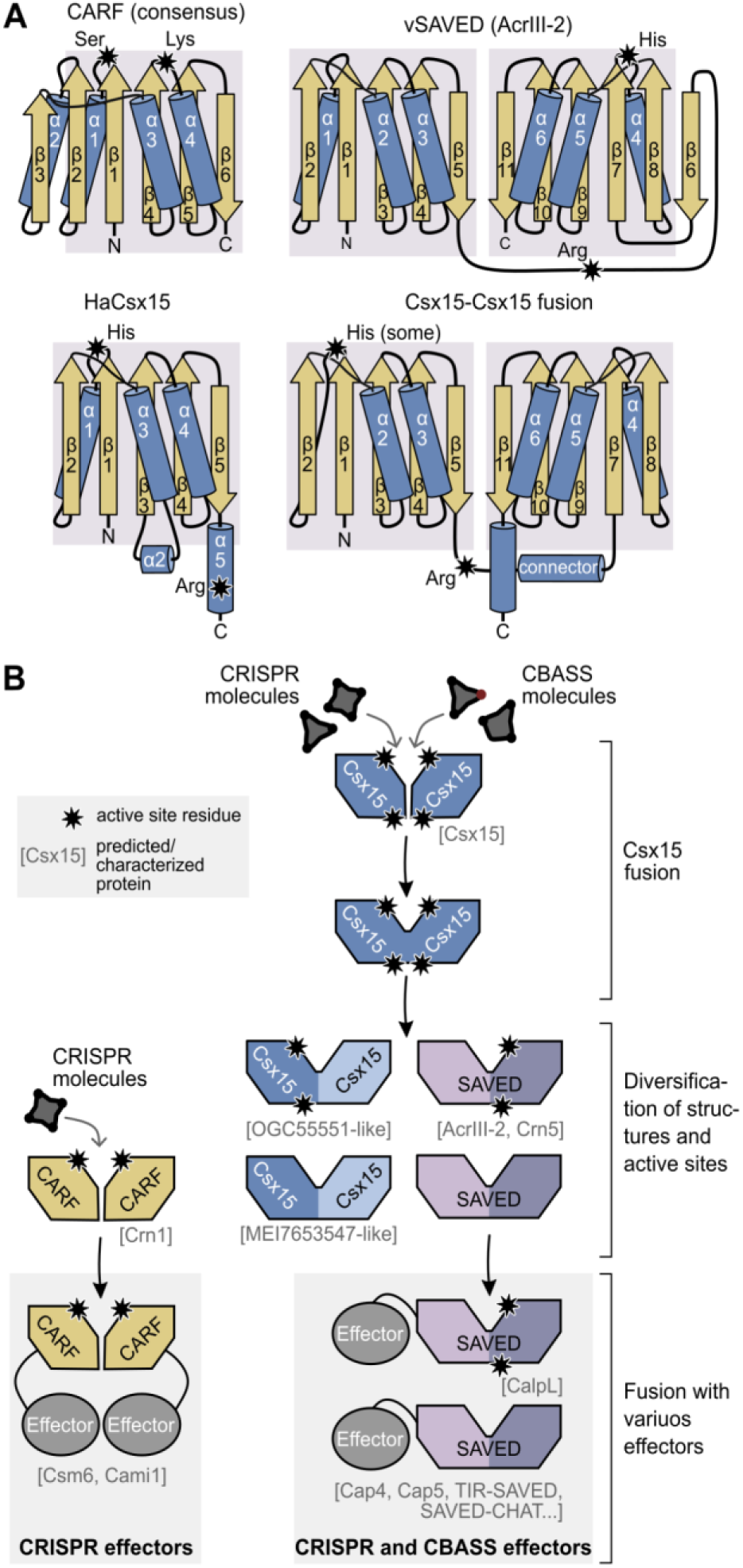
Evolutionary relationship between Csx15 and SAVED domain. (A) Structural topology of CARF, vSAVED (PDB 9R8S), HaCsx15, and Csx15-Csx15 fusion proteins. Compared with canonical CARF domains, Csx15 and each SAVED subdomain lack one α–β motif. Stars indicate catalytic residues (predicted in the Csx15-Csx15 fusion protein). (B) Proposed evolutionary relationship between Csx15 and SAVED domains. Canonical CARF proteins (left) binds symmetric cA_4_ or cA_6_ and function as homodimers with two identical active sites located on one side of the protein. In contrast, dimeric Csx15 (right) which bind both symmetric and asymmetric cyclic nucleotides and assemble into filaments similarly to SAVED proteins, are homologous to monomeric SAVED domains and contain two identical active sites positioned on opposite sides of the protein. Cyclic nucleotide sensor domains are shown as hexagons (CARF, Csx15) or nonagons (Csx15-Csx15, SAVED); and effector domains as ovals. Active sites are marked by stars. A characteristic example of each protein is indicated.

### Csx15 distribution and functional implications

Our genomic analyses indicate that Csx15 proteins are strongly associated with type III CRISPR-Cas systems. We found that most genomes and metagenomes containing a Csx15 homolog also encode a Cas10 cyclase of cA_n_ signaling. Furthermore, Csx15 and Cas10 frequently occur in close genomic proximity, providing strong evidence for a functional relationship between these proteins. This genomic association, together with the ring nuclease activity characterized here, supports a model in which Csx15 proteins act as specialized regulators or terminators of Cas10-derived cA_n_ signaling.

In contrast, the association between Csx15 proteins and CBASS systems is considerably weaker. Although Csx15 and CBASS cyclases occasionally co-occur within the same organism, such cases are substantially less frequent than the co-occurrence of Csx15 with Cas10. Consistent with this observation, CBASS cyclases were only detected in some Herpetosiphon genomes (*Herpetosiphon geysericola* and *Herpetosiphon llansteffanensis*), but were absent from the *Caldanaerobacter subterraneus* and *Herpetosiphon aurantiacus*, the source organisms of the Csx15 proteins in this study. Therefore, while our biochemical data demonstrate that Csx15 can degrade CBASS-derived cyclic nucleotides, the physiological significance of this activity remains unclear. Whether Csx15 participates in crosstalk between CRISPR-Cas and CBASS signaling pathways remains an important question for future investigation.

Interestingly, in contrast to previous reports^33^, we identified Csx15 homologs in viral genomes. However, these occurrences are rare and are predominantly restricted to prophage regions within the IMG/VR database. This limited distribution suggests that Csx15 is not commonly employed by viruses and may reflect its enzymatic properties. Csx15 rapidly cleaves cyclic nucleotides but exhibits slow turnover (at least for double-filament forming HaCsx15), likely due to the requirement for filament disassembly and reassembly after catalysis. While such kinetics may be well suited to bacterial CRISPR-Cas systems by facilitating clearance of signaling molecules after infection, viruses would likely favor enzymes with faster turnover for efficient immune suppression.

Despite this apparent limitation, the structural plasticity of Csx15 offers functional advantages. Although the symmetric Csx15 dimer is adapted to bind cA_4_, it can also accommodate asymmetric ligands such as cA_3_ and 3′3′3′-cAAG. This flexibility raises the possibility that viral Csx15 homologs function as broad-spectrum ring nucleases targeting multiple signaling pathways, including CRISPR-Cas and CBASS systems. Such adaptability mirrors the evolutionary strategy of viral 2H phosphodiesterases, which cleave diverse oligonucleotides and enable evasion of multiple bacterial defense systems, including type III CRISPR-Cas, CBASS, Panoptes and nucleotide-based signaling pathways involved in cellular regulation^46^.

### Csx15 fusions with other catalytic and non-catalytic domains

A number of Csx15 groups were found to be N-terminally fused to other catalytic or regulatory domains including Schlafen, Csx1, EAD, HTH domains and transmembrane helices (TMHs). Mammalian Schlafen (Slfn) domains are known to have endoribonuclease function that suppresses protein synthesis by cleaving tRNA and rRNA^47^. Slfn domains have a conserved active site signature, Ex_4_(D/E)xK motif that is essential for tRNA cleavage^47^. We found that prokaryotic Slfn-Csx15 fusion proteins retained Slfn active site suggesting they are able to target tRNAs and/or rRNAs (Figure S9). Csx1 are multidomain proteins that bind cA_n_ by the N-terminal CARF domain and this binding signal activates the C-terminal HEPN ribonuclease^48^. CARF domains of some Csx1 additionally have ring nuclease activity^19^. Csx1-Csx15 fusion proteins lack a canonical ring nuclease motif in Csx1 CARF domains, have intact HEPN active site and the Csx15 active site (Figure S10). Thus, in the case of Csx1-Csx15 fusion the Csx15 region seems to serve as a cA_n_ signal quenching element. Other domains fused with Csx15 do not have catalytic sites, but instead are likely to be involved in binding. Effector-associated domains (EADs) are predicted to function as adaptor domains mediating protein-protein interactions^49^, HTH domains typically function as DNA-binding domains, whereas TMH may be used either to tether proteins to the membrane or help to disrupt the membrane.

## MATERIALS AND METHODS

### Sequence search and analysis methods

Csx15 homologs were collected using PSI-BLAST^50^ search (5 iterations, E-value=0.005 inclusion threshold) with the *Caldanaerobacter subterraneus* protein (CsCsx15; WP_011026657)^32^ as a query against clustered non-redundant NCBI protein sequence database (ClusteredNR). Viral Csx15 homologs specifically were searched in the NCBI database of metagenomics proteins (env_nr) and the IMG/VR database^51^ with several Csx15 queries using BLAST^50^. Within the IMG/VR search results, only high-confidence viral metagenomes of high and medium quality level of completeness were considered.

Protein grouping by sequence similarity was done by clustering with CLANS^52^. Sequence alignment was performed with MAFFT^53^ using an accuracy-oriented mode (L-INS-i).

Initial identification of conserved protein domains was done using sequence search against the NCBI conserved domain database (CDD)^54^ search. Domain composition of sequences/regions lacking confident CDD assignments were further explored with HHsearch^55^. In addition, domain assignments were validated using AlphaFold models for representatives of specific domain architectures. Prediction of transmembrane helices in protein sequences was performed with TMHMM - 2.0^56^.

Identification and typing of CRISPR-Cas systems and associated components was performed using CRISPRCasTyper^57^ and PADLOC^58^.

For the analysis of co-occurrence of Csx15, Cas10, CdnD, CdnE, and DncV, 1204 (meta)genomes with at least 2000 open reading frames per genome containing an annotated open reading frame for Csx15 were obtained from NCBI. Phylogeny was inferred using ETE3^59^. hmmsearch^60^ was used to query for the proteins of interest. To obtain the query profiles, Csx15, Cas10, CdnD, CdnE, and DncV, alignments were generated using blastp,^61^ curated to remove fusion proteins and truncated sequences, and then converted to .hmm search profiles using hmmbuild^61^. Species counts were plotted on the ETE3 generated phylogenetic tree using ggtree^62^. Gene neighbourhoods were analyzed using custom python scripts.

### Structure modeling and analysis

Structural models of sequence representatives were either taken from the AlphaFold database (AFDB)^63^ or modeled using the AlphaFold3^64^ web server. Structure search and comparison was performed with either Foldseek^65^ or DALI^66^. Model visualization and interactive analysis was done using ChimeraX^67^.

### Cloning and mutagenesis

The coding sequences of the *Herpetosiphon aurantiacus* DSM 785 and *Caldanaerobacter subterraneus subsp. tengcongensis* strain MB4 Csx15 proteins, were obtained from the NCBI reference genome sequences CP000875.1 (protein ABX03341.1) and NC_003869.1 (protein WP_011026657.1), respectively. Both gene sequences of the proteins have been optimized for expression in *E. coli*. The *E. coli* DH5α strain was used for cloning and plasmid amplification. Table S4 contains a list of all the vectors constructed in this study.

Synthetic plasmids encoding *Csx15* (fused to TEV-cleavable His_6_-StrepII-His_6_ N-tag) genes were purchased from Twist Bioscience. Fragments containing *Csx15* genes were recloned into the pMBP plasmid to obtain pMBP-HaCsx15 and pMBP-CsCsx15 plasmids. To generate plasmids encoding mutant versions of HaCsx15 and CsCsx15, site-directed mutagenesis was used with primers carrying the desired mutations. The Gibson assembly strategy was used to clone AvCami1-S11A alone or together with HaCsx15 into pETDuet-1 vector (Novogene) to generate plasmids for *in vivo* experiments.

### Expression and purification of proteins

N-tagged WT HaCsx15 was expressed in *E. coli* ArcticExpress (DE3), whereas mutant HaCsx15 variants and both WT and mutant CsCsx15 variants were expressed in BL21 (DE3). Bacteria were grown at 37°C in LB medium supplemented with ampicillin (100 µg/ml) to the mid-log phase. Expression was induced with 1 mM IPTG and cells were further cultured at 16°C overnight. Cells were harvested by centrifugation and disrupted by sonication. Soluble proteins were captured on the HisTrap HP column (Cytiva) using chromatography buffer A1 (20 mM Tris-HCl (pH 7.4), 0.5 M NaCl, 7 mM 2-mercaptoethanol) and a 40-500 mM imidazole gradient. The eluted fractions were pooled and imidazole was removed by loading the protein on HiPrep 26/10 Desalting column (Cytiva). To remove the affinity tag from the protein, the eluted fractions were pooled and incubated with TEV protease^68^ (at a ratio 1:50 (w/w) protease:target protein) and 2 mM DTT at 4°C overnight. After the TEV cleavage reaction, the cleaved affinity tag and TEV protease were removed by loading the sample again on a HisTrap HP column (Cytiva). Peak fractions of the eluted sample were dialyzed against storage buffer (10 mM Tris-HCl (pH 7.4), 300 mM NaCl, 1 mM DTT, 0.1 mM EDTA and 50% (v/v) glycerol), and stored at −20°C. The purified proteins were analyzed by SDS-PAGE.

### cA_n_ radiolabeling and purification

α^32^P-labeled cA_3_ was prepared by mixing 0.2 µM StCsm complex with 0.2 µM S3/1 target RNA (5’-GGGCGGCAAAUUGAGGAGGUGGAAUAAGUGAACAGAAUUAAACAGUUACGAAAAACAAAAGGUGGCGA-3’)^39^, 10 mM CoCl_2_, 0.2 mM ATP and 1 µM α^32^P-ATP in the reaction buffer Y (33 mM Tris-acetate (pH 7.9), 66 mM K-acetate, 1 mM DTT) and incubating at 37°C for 4 hours. α^32^P-labeled cA_4_ was prepared by mixing 0.15 µM CCaCsm complex with 10 µM target RNA (5’-GGGUGAAGAGCAAUGAGCUCUCGAGGUGCGAUAUCGCUCUUCCCAGUGUA-3’)^18^ prepared by *in vitro* transcription, 1 mM Mg(CH_3_COO)_2_, 0.2 mM ATP and 1 µM α^32^P-ATP in the reaction buffer H and incubating at 37°C for 4 hours. Similarly, α^32^P-labeled cA_5_ and cA_6_ was prepared by mixing 0.2 µM StCsm complex with 10 µM S3/1 target RNA. Synthesis products were separated by denaturing PAGE (34.5% gel, 19:1 acrylamide: bisacrylamide ratio, 6 M urea in 0.5× TBE). Individual cA_n_ species were excised from the gel and purified by extraction in 1 mM EDTA with shaking at 1000 rpm overnight at 4 °C, followed by ethanol precipitation.

### Ring nuclease activity reactions

Cyclic oligonucleotide hydrolysis assays were performed in reaction buffer Y for HaCsx15 or reaction buffer C (20 mM sodium cacodylate (pH 6.5), 60 mM NaCl, 1 mM DTT) for CsCsx15 reactions, supplemented with 0.1 mg/ml BSA (Thermo Fisher Scientific), and containing 10 µM synthetic cA_3_, cA_4_, cA_5_, cA_6_, 2′3′3′-cAAA, 3′3′3′-cAAG, 3′3′-c-di-AMP, 3′3′-cGAMP, A_2_p, or A_2_>p (Biolog). Reactions were initiated by the addition of 13 µM protein (WT HaCsx15 or CsCsx15) and incubated at 37°C for the indicated times. Reactions were terminated by adding Proteinase K to a final concentration of 1 mg/ml (Thermo Fisher Scientific) followed by incubation at 55°C for 10 min. Reaction products were analyzed by HPLC-MS.

To directly visualize cA_n_ hydrolysis, single-turnover reactions were performed in reaction buffer Y for HaCsx15 or reaction buffer C for CsCsx15 reactions, supplemented with 0.1 mg/ml BSA (Thermo Fisher Scientific) and containing 10 nM radiolabeled α^32^P-cA_3_, α^32^P-cA_4_, α^32^P-cA_5_, or α^32^P-cA_6_ together with 490 nM of the corresponding unlabeled cA_n_ (Biolog). Reactions were initiated by addition of 5 µM protein (WT or mutant Csx15) and incubated at 37°C. Multiple-turnover reactions contained 10 nM radiolabeled α^32^P-cA_3_, α^32^P-cA_4_, α^32^P-cA_5_, or α^32^P-cA_6_ together with 490 nM of the corresponding unlabeled cA_n_ (Biolog), and were initiated by addition of 50 nM protein. At the indicated time points, 5 µl aliquots were withdrawn and quenched by mixing with an equal volume of 2× RNA Loading Dye (Thermo Fisher Scientific). Reaction products were analyzed by denaturing PAGE (30% gels, 19:1 acrylamide:bis-acrylamide ratio, 6 M urea in 0.5× TBE) and visualized by autoradiography.

### HPLC-MS analysis

To analyze the samples, electrospray ionization mass spectrometry (ESI-MS) was performed in negative mode using an integrated HPLC/ESI-MS system (1290 Infinity II, Agilent Technologies/Q-TOF 6575 Revident, Agilent Technologies) equipped with a Supelco Discovery®HS C18 column (7.5 cm × 2.1 mm, 3 µm). Elution was performed with a linear gradient of solvents A (5 mM ammonium acetate in water, pH 7.0) and B (5 mM ammonium acetate in methanol, pH 7.0) at a flow rate of 0.3 ml/min at 30°C as follows: 0–3 min, 0% B; 3–19 min, 0-12% B; 19–21 min, 12-90% B, 21–25 min 90% B. Ionization capillary voltage was set to 3500 V, fragmentor to - 175V. Molecules were annotated based on accurate mass and retention time from synthesized standards.

### SEC-MALS analysis

SEC-MALS analysis was performed on an HPLC system (Waters Breeze) combined with a miniDAWN MALS detector (Wyatt) and an Optilab refractive index detector (Wyatt). HaCsx15 WT or mutant protein samples were diluted with reaction buffer H2 (20 mM HEPES (pH 7.4), 60 mM KCl, 1 mM DTT), CsCsx15 WT samples were diluted with reaction buffer C to a final concentration of 0.5 mg/ml. For oligomerization studies, WT protein samples were supplemented with a five-fold molar excess of F_4_cA_4_ (Biolog), calculated relative to the molar amount of Csx15 dimers. 200 µL of diluted samples were loaded to a Superdex 200 Increase 10/300 GL column (Cytiva) equilibrated with corresponding buffer. Data acquisition and evaluation were performed using ASTRA 7 software (Wyatt).

### Mass photometry analysis

The WT HaCsx15 sample was diluted in PBS (phosphate-buffered saline) buffer (10 mM Na_2_HPO_4_, 1.8 mM KH_2_PO_4,_ 137 mM NaCl, 2.7 mM KCl, pH 7.4), while the WT CstCsx15 sample was diluted in reaction buffer C with (pH 7.4), respectively. Both buffers were purified by ultrafiltration using 3 kDa Amicon columns.

The WT HaCsx15 and WT CsCsx15 proteins, which were used to measure weight distribution analysis Figure S4A and WT HaCsx15 concentration-dependent filament formation Figure S4B, were diluted to 1 µM. To measure weight distribution, the proteins were diluted to 100 nM and analyzed. In the latter experiment, the cA_4_ ligand was diluted to 20 µM in PBS buffer, and then mixed with the protein to form a 500 nM complex at a 1:20 molar ratio of protein to ligand. The mixture was then quickly diluted to 250 nM, 100 nM and 50 nM, while preserving the 1:20 molar ratio. For apo-state controls in concentration-dependent filament formation experiments, the proteins were diluted only with buffer. Time zero was defined as the moment of mixing the protein and cA_4_ ligand (or buffer for the controls), and mass photometry measurements were taken after 40 minutes of incubation.

The measurements were performed at room temperature using a Refeyn TWO MP apparatus (Refeyn Ltd., Oxford, UK). Each measurement took 60 seconds. The results were analyzed and the figures were created using Refeyn Discovery software (DiscoverMP 2023 R1.2).

### Dynamic light scattering

A 5 μl sample was prepared by mixing 12.5 μM HaCsx15, CsCsx15 WT or mutant protein samples with 25 µM cA_3_, cA_4_, cA_5_, cA_6,_ 2′3′3′-cAAA, 3′3′3′-cAAG, A_2_p, A_2_>p (Biolog) or buffer control. Reaction buffer Y was used for HaCsx15 samples, and reaction buffer C was used for CsCsx15 samples. The sample was loaded into a 2 μl Quartz Cuvette (Malvern Panalytical), and measurements were carried out at 22 °C with a Zetasizer μV photometer (Malvern Panalytical) using Zetasizer Software (v6.20). Measurements were carried out in 60 s runs containing 6 × 10 s runs with 10 measurements in total. For time-course experiments, 150 measurements in total were performed.

### HaCsx15-F_4_cA_4_ crystallization

The HaCsx15 complex was crystallized at 18°C using the sitting-drop vapor diffusion method. The HaCsx15 reservoir protein was concentrated and re-buffered in reaction buffer H1 (20 mM HEPES (pH 7.4), 150 mM KCl and 1 mM DTT), and then mixed with the F_4_cA_4_ ligand at a molar ratio of 1:2 to reach a final concentration of 21 mg/ml. The mixture was then incubated at 4°C for 30 minutes before being used to set up crystallization trials. Crystallization drops were set up using a Crystal Phoenix robot (Art Robbins Instruments). The drops consisted of 0.3 µl of reservoir buffer and 0.3 µl of the mixture. After three months, crystals were obtained for the Morpheus G5 condition: 100 mM MOPS/HEPES-NaOH (pH 7.5), 20 mM sodium formate, 20 mM ammonium acetate, 20 mM trisodium citrate, 20 mM sodium potassium L-tartrate, 20 mM sodium oxamate, 10% w/v PEG 20,000 and 20% v/v PEG MME 550. As the Morpheus G5 condition was already cryo-protective, the crystals were fished out of the drop directly and flash frozen in liquid nitrogen.

### Data collection and refinement

The diffraction data for the HaCsx15-F_4_cA_4_ crystals were collected at the P14.1 beamline at the BESSY II synchrotron in Berlin. The diffraction data were processed using XDS^69^. The data collection statistics are presented in Table 1. The HaCsx15-F_4_cA_4_ crystals belonged to the P3(1)21 space group and contained one HaCsx15 dimer and one F_4_cA_4_ molecule in the asymmetric unit. The structure was solved by molecular replacement using PHASER^70^, with the dimer search model generated by AlphaFold3^64^ being subsequently rebuilt in Autobuild to improve its fit with the electron density. Manual model building was performed in COOT^71^. Refinement was performed using PHENIX^72^. The diffraction data parameters and final refinement statistics are listed in Table 1.

### Cryo-EM sample preparation

The HaCsx15 sample with F_4_cA_4_ was prepared by mixing 68.2 µM HaCsx15 WT (in terms of dimer, 2 mg/mL) and 64 µM of F_4_cA_4_ in the reaction buffer H1 supplemented with 7 mM 2-mercaptoethanol. The CsCsx15 samples were prepared by mixing 78 µM CsCsx15 WT (in terms of dimer, 2.3 mg/mL) and 250 µM cA_4_ or 250 µM 3′3′3′-cAAG in the reaction buffer C. The HaCsx15 sample was incubated for 60 min at 4 °C before grid preparation, whereas CsCsx15 samples were used immediately after mixing. The samples were applied on freshly glow-discharged (20 mA, 45 s in a GloQube Plus device (Quorum Technologies)) Quantifoil Cu 300 mesh R1.2/1.3 grids using a blotting time of 5 s under 95% humidity conditions at 4° C and plunge-frozen into liquid ethane using Vitrobot Mark IV device (FEI).

### Cryogenic-electron microscopy data collection, processing, model building, and refinement

Cryogenic-electron microscopy (cryo-EM) data collection on a Glacios microscope (Thermo Fisher Scientific) and processing in CryoSPARC^73^ for the HaCsx15-F_4_cA_4_ and CsCsx15-A_2_>P structures are summarized in Figure S6. The initial models for both proteins were obtained using AlphaFold3^64^. Multiple copies of the model were fitted into the respective cryo-EM maps using ChimeraX (v.1.7)^67^. Protein rebuilding and manual building of oligoadenylates were performed using Coot (v.0.8.9)^71^. Model refinement was performed using phenix.real_space_refine (v. 1.20.1_4487)^72^; refinement statistics are summarized in Table 2. Structure overlays and generation of structural images were performed using ChimeraX (v.1.7)^67^.

Cryo-EM data collection and initial analysis for the CsCsx15 complex with 3′3′3′-cAAG followed the same procedures as described for the CsCsx15 complex with A_2_>p (Table 2 and Figure S6): 2,055,030 particles (256 px box size, pixel size 1.1 Å) were extracted from 1273 accepted exposures. However, due to the relatively small and irregular size of the resultant filament-like particles, further analysis was limited to 2D classification.

### *E. coli* survivability assays

*E. coli* ER2566 cells were co-transformed with pCsm_dNucleases, pCRISPR and pTarget plasmids in combination with (i) pAvCami1-S11A, (ii) pAvCami1-S11A_HaCsx15 or (iii) pAvCami1-S11A_HaCsx15-H8A (Table S4), plated on LB media agar plates, supplemented with streptomycin (50 µg/ml), carbenicillin (50 µg/ml), chloramphenicol (25 µg/ml), kanamycin (50 µg/ml) and 1% glucose and grown overnight at 37°C. A single colony of each variant was grown overnight at 37°C in liquid LB supplemented with streptomycin (50 µg/ml), carbenicillin (50 µg/ml), chloramphenicol (25 µg/ml), kanamycin (50 µg/ml) and 1% glucose. The overnight cultures were diluted 1:200 in the fresh LB containing streptomycin (50 µg/ml), carbenicillin (50 µg/ml), chloramphenicol (25 µg/ml) and kanamycin (50 µg/ml) and incubated in a 96-well plate for 1 hour at 37°C with shaking. Then 0.1 mM IPTG was added to induce the protein and crRNA expression. After induction bacterial growth was measured by recording optical density at 600 nm (OD_600_) every 10 minutes using the CLARIOstar Plus Microplate Reader (BMG Labtech). The mean with standard deviation of five biological replicates is presented for each experiment.

## RESOURCE AVAILABILITY

### Materials availability

Plasmids generated in this study are available upon request from the lead contact with a completed Materials Transfer Agreement.

### Data availability

The crystal diffraction data (coordinates and structure factors) were deposited in the Protein Data Bank under accession code 30JQ (HaCsx15 complex with F_4_cA_4_). The EM densities and model coordinates have been deposited in the Electron Microscopy Data Bank and Protein Data Bank under the accession codes EMD-56908/PDB:28WJ (HaCsx15 complex with F_4_cA_4_), and EMD-57705/PDB:30FQ (CsCsx15 complex with A_2_>p).

## ACKNOWLEDGMENTS

We thank Irmantas Mogila for his help with the preparation of Figure 1A from the CLANS analysis, Kryzsztof Skowronek for help with mass photometry, and Virginijus Siksnys for discussion. This work was supported by the Research Council of Lithuania (LMTLT) (grant S-LL-24-6 to G.T.) and by the National Science Centre of Poland (grant 2024/52/L/NZ1/00249 to M.B.). Core facility (RRID:SCR_021630) support was funded by the IIMCB IN-MOL-CELL Infrastructure (RRID: SCR_021630) funded by the European Union – Next Generation EU under National Recovery and Resilience Plan. IN-MOLCELL Infrastructure was also funded by the European Union under Horizon Europe (Project 101059801 - RACE) and by RACE-PRIME project carried out within the IRAP programme of the Foundation for Polish Science co-financed by the European Union under the European Funds for Smart Economy 2021-2027 (FENG). We are grateful for access to the BESSY (Berlin) beamline MX 14.1.

## DECLARATION OF GENERATIVE AI AND AI-ASSISTED TECHNOLOGIES IN THE WRITING PROCESS

During the preparation of this work, the authors used DeepL/Write or ChatGPT (OpenAI) in order to improve the readability and language of the manuscript. After using these tools, the authors reviewed and edited the content as needed and take full responsibility for the content of the published article.

