## Supplemental Data for "Csx15 is a broad specificity ring nuclease bridging CRISPR-Cas and CBASS cyclic nucleotide signaling systems"

### SUPPLEMENTARY FIGURES & TABLES

**Figure S1. Multiple sequence alignment of the Csx15-Csx15 fusion proteins.** Related to Figure 1.

**Figure S2. Csx15 proteins used in this study.** Related to Figure 1.

**Figure S3. Ring nuclease activity of Csx15. Related to Figure 1.**

**Figure S4. Substrate-induced oligomerization of Csx15.** Related to Figures 1 and 2.

**Figure S5. HaCsx15 structural similarity to CARF- and SAVED-effectors.** Related to Figure 2.

**Figure S6. Cryo-EM single particle analysis workflow.** Related to Figure 3.

**Figure S7. Oligomeric states of putative active site mutants of HaCsx15.** Related to Figures 2, 3 and 4.

**Figure S8**. **Multiple sequence alignment of putative viral Csx15 homologs.** Related to Figure 7.

**Figure S9**. **Multiple sequence alignment of the Schlafen-Csx15 fusion proteins.** Related to Figure 1.

**Figure S10. Multiple sequence alignment of the Csx15 fusion with Csx1.** Related to Figure 1.

**Table S1. Information for Csx15 proteins analyzed in this work (separate file).** Related to Figure 1.

**Table S2. Domain architectures of the central Csx15 cluster (separate file).** Related to Figure 1.

**Table S3. Csx15 proteins analyzed in co-occurrence with CRISPR-Cas and CBASS signaling systems (separate file).** Related to Figure 6.

**Table S4. Plasmids used in this study.** Related to Figures 1, 2, 4 and 5.

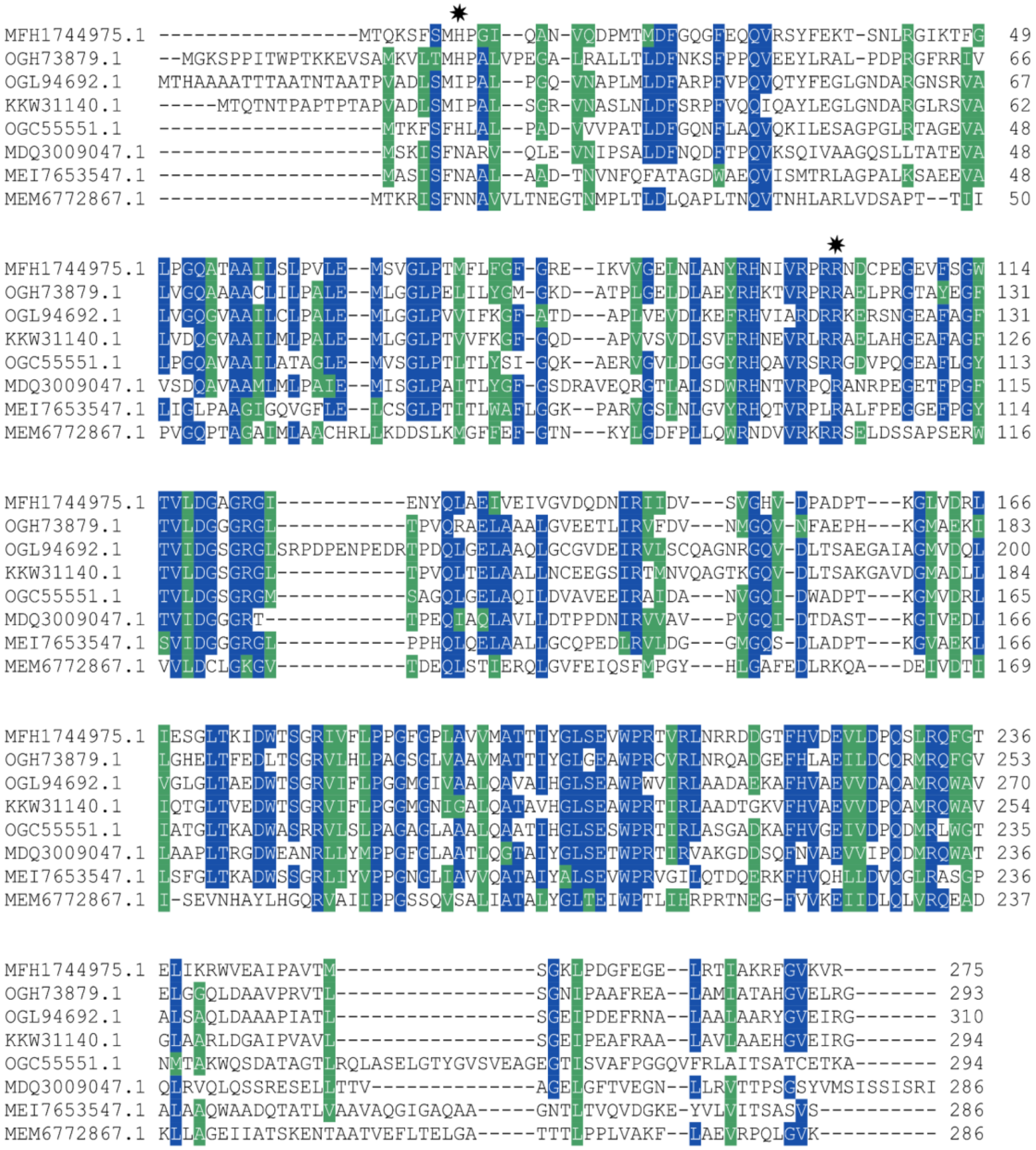


**Figure S1. Multiple sequence alignment of the Csx15-Csx15 fusion proteins. Related to Figure 1.** Putative active sites in the Csx15-Csx15 proteins are indicated with black stars above the alignment.

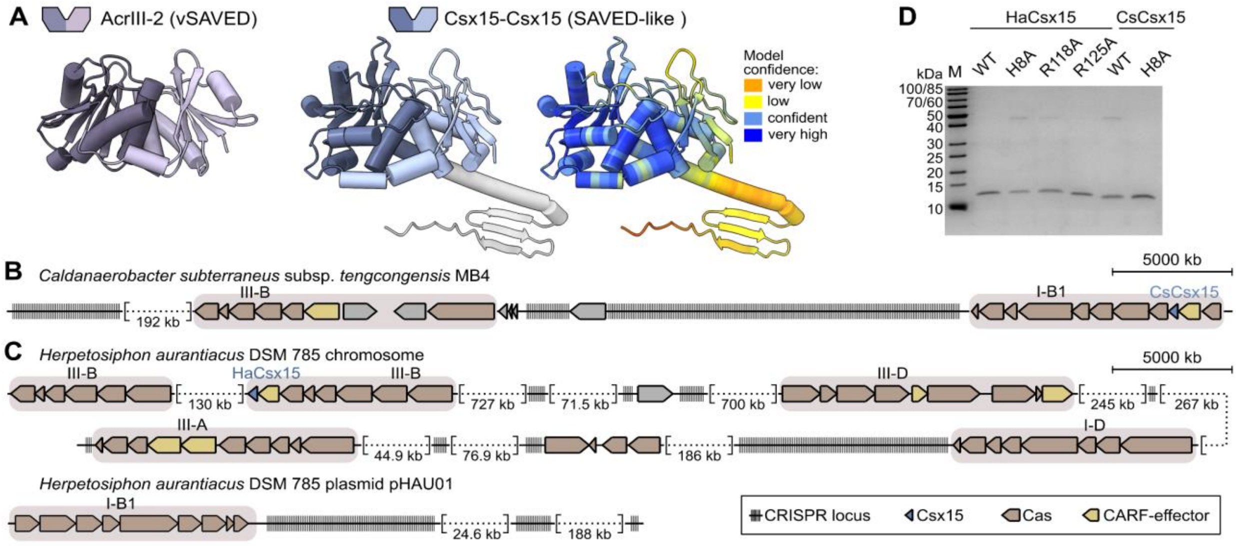


**Figure S2. Csx15 proteins used in this study. Related to Figure 1.**

(A) Crystal structure of a viral vSAVED (AcrIII-2) (left) (PDB: 9R8S)[1] and AlphaFold3 model of a SAVED-like Csx15-Csx15 (NCBI: OGC55551) (right).

(B) Detailed schematic representation of the genomic neighborhoods of CsCsx15 selected for characterization. The complete genome of *C. subterraneus* subsp. *tengcongensis* MB4 (NCBI: AE008691.1) contains the Csx15 protein (AAM25784.1) in close proximity to the Type I-B CRISPR-Cas system and the neighbouring CARF-HTH protein (AAM25785.1). The Type III-B system encodes the Csx1 protein (AAM25756.1).

(C) Detailed schematic representation of the genomic neighborhoods of HaCsx15 selected for characterization.The complete chromosome sequence of *Herpetosiphon aurantiacus* DSM 785 (NCBI: CP000875.1) contains several CRISPR-Cas systems. HaCsx15 is encoded adjacent to the Type III-B system next to the Can2/Card1 homolog (ABX03342.1). Additionally, two other CARF proteins (ABX05118.1, ABX05119.1) are encoded next to the Type III-A. A CARF protein (ABX04565.1) is encoded in the vicinity of the Type III-D system. *H. aurantiacus* has two plasmids, pHAU01 (CP000876.1) and pHAU02 (CP000877.1). The larger plasmid (pHAU01) encodes a complete Type-IB CRISPR-Cas system, but no CARF proteins are encoded in its vicinity. The smaller plasmid does not have any CRISPR-Cas systems.

(D) SDS-PAGE of Csx15 proteins used in this study.

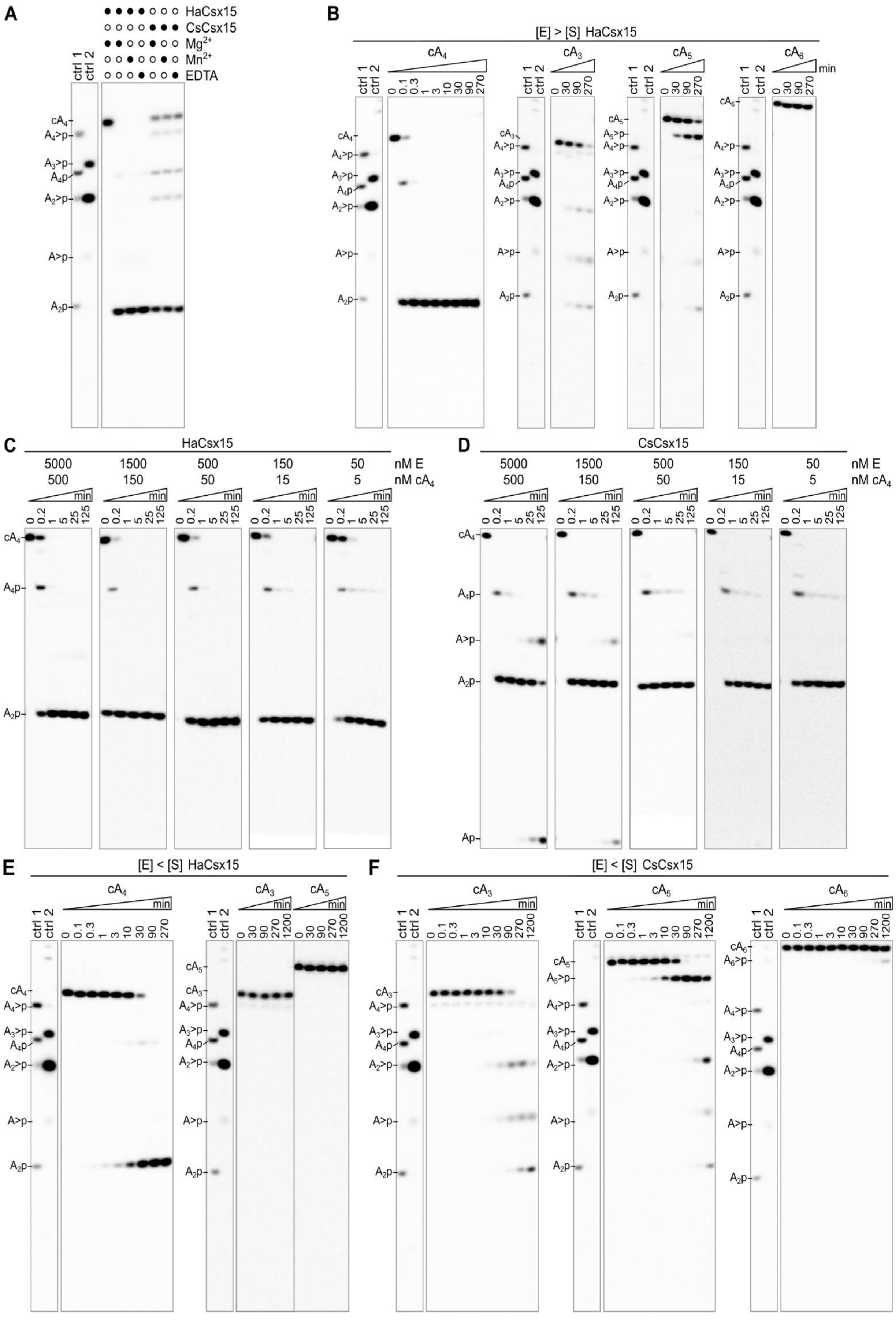

A
HaCsx15
CsCsx15
Mg2+
Mn2+
EDTA
ctrl 1
ctrl 2
cA4
A4>p
A3>p
A4p
A2>p
A>p
A2p
B
[E] > [S] HaCsx15
cA4
cA3
cA5
cA6
min
0 0.1 0.3 1 3 10 30 90 270
0 30 90 270
A5>p
C
HaCsx15
D
CsCsx15
5000 1500 500 150 50 nM E
500 150 50 15 5 nM cA4
0 0.2 1 5 25 125
Ap
E
[E] < [S] HaCsx15
0 30 90 270 1200
F
[E] < [S] CsCsx15
0 0.1 0.3 1 3 10 30 90 270 1200
A6>p

**Figure S3. Ring nuclease activity of Csx15. Related to Figure 1.**

(A) Cleavage of $^{32}$P-labeled $cA_4$ by WT Csx15 in the presence or absence of divalent metal ions under multiple-turnover conditions (60 min for HaCsx15; 5 min for CsCsx15). Reactions contained 10 mM $Mg(CH_3COO)_2$, 1 mM $MnCl_2$, or 1 mM EDTA.

(B) Cleavage of $^{32}$P-labeled $cA_4$, $cA_3$, $cA_5$ and $cA_6$ by WT HaCsx15 under single-turnover conditions.

(C, D) Cleavage of 5-500 nM $^{32}$P-labeled $cA_4$ by WT HaCsx15 (C) or CsCsx15 (D) under single-turnover conditions.

(E) Cleavage of $^{32}$P-labeled $cA_4$ and $cA_3$ by WT HaCsx15 under multiple-turnover conditions.

(F) Cleavage of $^{32}$P-labeled $cA_3$, $cA_5$ and $cA_6$ by WT CsCsx15 under multiple-turnover conditions.

Unless indicated otherwise (in C, D), reactions contained 500 nM $^{32}$P-labeled $cA_n$ and 5 µM (single-turnover) or 50 nM (multiple-turnover) WT HaCsx15 or CsCsx15 and were analyzed by PAGE. $cA_4$ cleavage by CCaCalpL (ctrl 1) and $cA_6$ cleavage by StCsm6 (ctrl 2) served as markers for identification of Csx15 reaction intermediates and products.

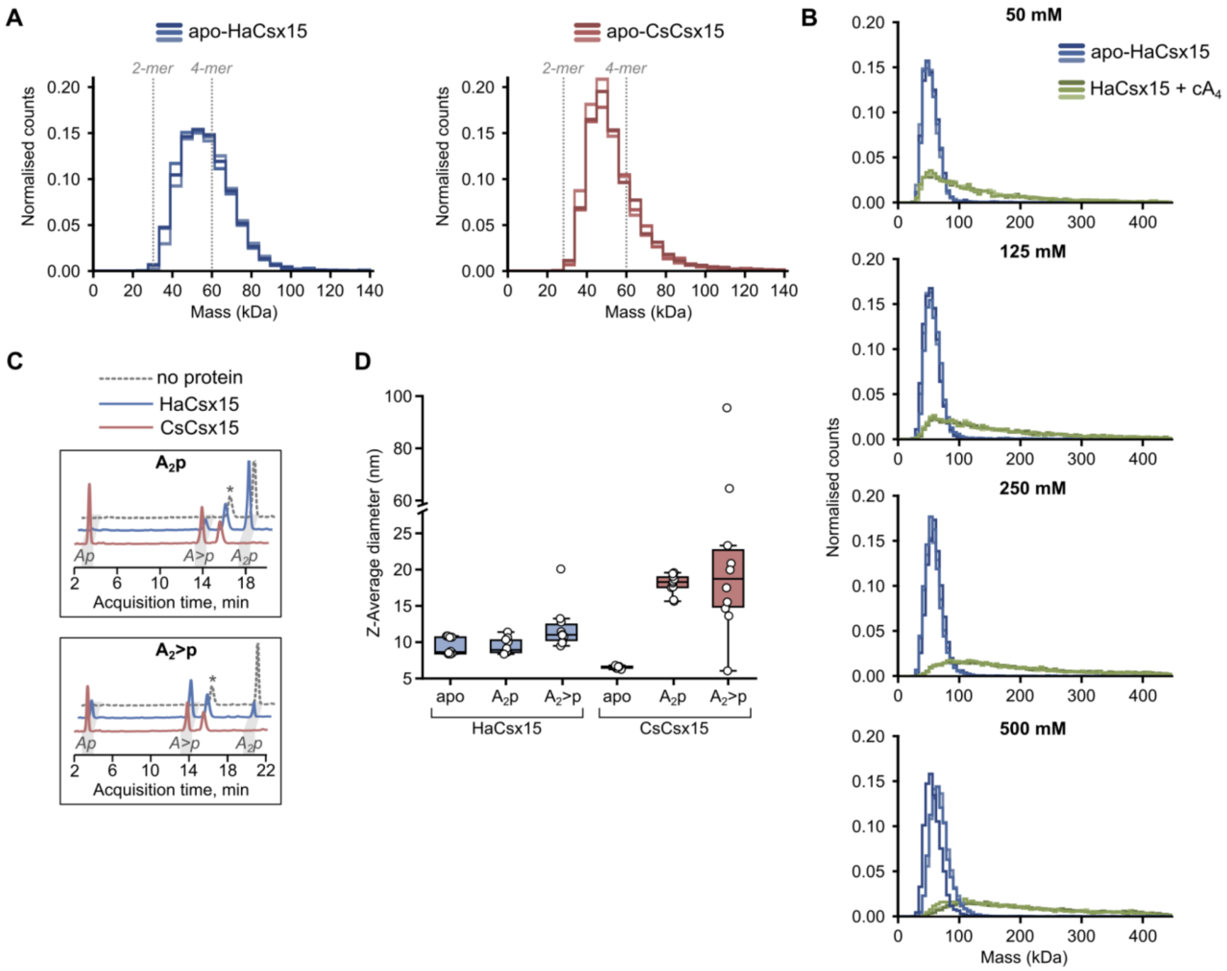


**Figure S4. Substrate-induced oligomerization of Csx15. Related to Figures 1 and 2.**

(A) Mass photometry analysis of apo WT HaCsx15 (left) and CsCsx15 (right), showing their molecular weight distributions in solution.

(B) Concentration-dependent oligomerization of WT HaCsx15 induced by $cA_4$ measured by mass photometry.

(C) Hydrolysis of the reaction intermediates $A_2p$ and $A_2>p$ by WT HaCsx15 and CsCsx15, analyzed by HPLC-MS. Reactions contained 10 μM di-nucleotide and 13 μM protein and were incubated for 2 h. Asterisk (*) denotes a buffer-derived peak.

(D) DLS analysis of $A_2p$- and $A_2>p$-induced oligomerization of WT HaCsx15 and CsCsx15. Data are shown as box plots (n = 10 measurements).

**A**

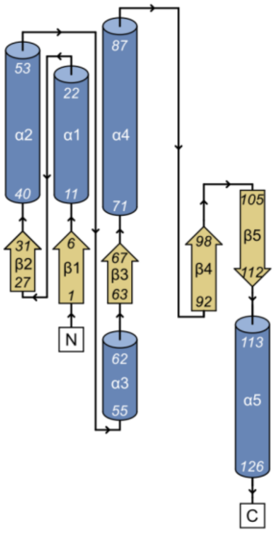


**B**

| Protein | Organism | PDB | DALI score | Domain | Signaling system | Signaling molecule | Ring nuclease |
|---|---|---|---|---|---|---|---|
| HaCsx15 | *Herpetosiphon aurantiacus* DSM 785 | 30JQ, 28WJ | - | Csx15 | type III CRISPR-Cas | $cA_3$, $cA_4$, 3'3'3'-cAAG, 2'3'3'-cAAA | yes (this study) |
| CsCsx15 | *Caldanaerobacter subterraneus* | 30FQ | - | Csx15 | type III CRISPR-Cas | $cA_3$, $cA_4$, $cA_5$, $cA_6$, 3'3'3'-cAAG, 2'3'3'-cAAA | yes (this study) |
| SAVED-CHAT | *Haliangium ochraceum* | 8TL0 | 8.0 | SAVED2 | type III CRISPR-Cas | $cA_3$ | no[2] |
| Cap4 | *Acinetobacter baumannii* | 6VM6 | 7.3 | SAVED1 | CBASS | 2'3'3'-cAAA | no[3] |
| Cap5 | *Pseudomonas syringae* | 8FM1 | 7.2 | SAVED1 | CBASS | 3'2'-c-di-AMP | no[4] |
| CalpL | *Candidatus* Cloacimonas acidaminovorans | 9EYI | 7.0 | SAVED4 | type III CRISPR-Cas | $cA_4$ | yes[5] |
| vSAVED | *Thermocrinis Great Boiling Spring virus* | 9R8S | 6.7 | SAVED4 | anti-CRISPR | $cA_4$ | yes[1] |
| TIR-SAVED | *Microbacterium terrae* | 7QQK | 6.5 | SAVED5 | CBASS | $cA_3$ | no[6] |
| Cap14 | *Bacillus thuringiensis Bt407* | 9OM7 | 6.4 | SAVED2 | CBASS | 2'3'-cGAMP | no[7] |
| Cap5 | *Lactococcus cremoris subsp. cremoris IBB477* | 7RWM | 5.9 | SAVED1 | CBASS | 3'2'-cGAMP | no[8] |
| Can2 | *Vibrio cholerae* | 1XMX | 5.7 | CARF4 | type III CRISPR-Cas | $cA_4$ | no[9] |
| Csa3 | *Saccharolobus solfataricus P2* | 2WTE | 5.7 | CARF3 | type III CRISPR-Cas | $cA_4$ | no[10] |
| Card1 | *Treponema succinifaciens* | 6XL1 | 5.7 | CARF4 | type III CRISPR-Cas | $cA_4$, $cA_6$ | no[11] |
| Crn1 (Sso2081) | *Saccharolobus solfataricus* | 7YGH | 5.7 | CARF_m13 | type III CRISPR-Cas | $cA_4$ | yes[12] |
| Cap4 | *Enterobacter cloacae* | 7YIB | 5.5 | SAVED1 | CBASS | 3'3'3'-cAAG | no[13] |
| Can2 | *Thermoanaerobacter brockii* | 8Q3Z | 5.5 | CARF4 | type III CRISPR-Cas | $cA_4$ | no[14] |
| CARF-ADA | *Thermoanaerobaculum aquaticum* | 9W1I | 5.7 | CARF5 | type III CRISPR-Cas | $cA_4$, $cA_6$ | yes ($cA_4$ only)[15] |
| Csx1 | *Sulfolobus islandicus* | 6QZQ | 5.4 | CARF2 | type III CRISPR-Cas | $cA_4$ | no[16] |
| Cap5 | *Deinococcus wulumuqiensis* | 8YSQ | 5.3 | SAVED1 | CBASS | 3'3'-cAA, 3'3'-cGG, 3'3'-cUU, 3'3'-cCC | no[17] |
| Cdn1 | *Psychrobacter lutiphocae* | 9U49 | 5.2 | CARF | type III CRISPR-Cas | $cA_3$, $cA_4$, $cA_6$ | no[18] |
| Crn1 (Sso1393) | *Saccharolobus solfataricus* | 3QYF | 5.0 | CARF7 | type III CRISPR-Cas | $cA_4$ | yes[19] |

**Figure S5. HaCsx15 structural similarity to CARF- and SAVED-effectors. Related to Figure 2.**
(A) Topology diagram of HaCsx15.
(B) Table listing proteins[2-19] with similar structures to HaCsx15. HaCsx15 monomer (PDB: 30JQ) was used as a query in DaliSearch[20].

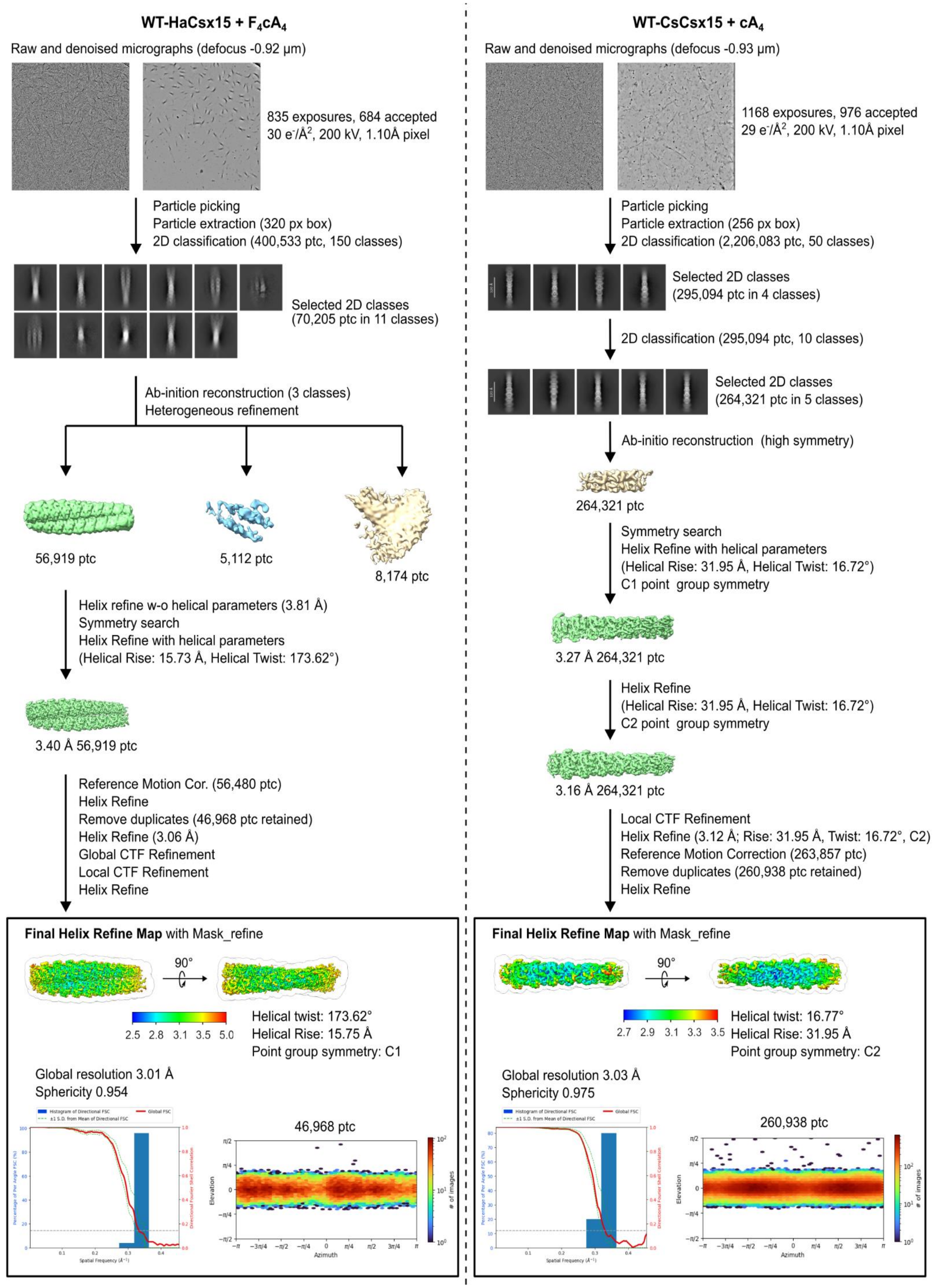


**Figure S6. Cryo-EM single particle analysis workflow. Related to Figure 3.**

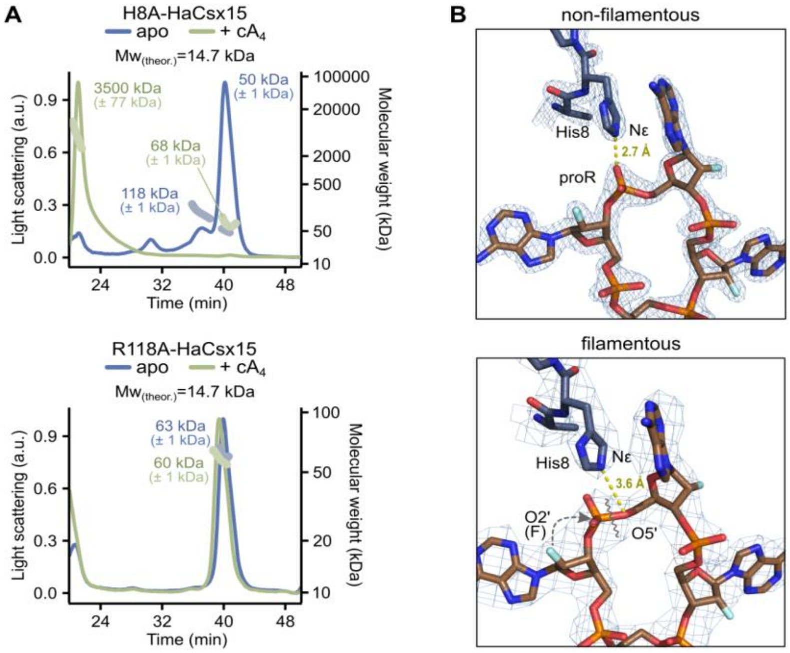


**Figure S7. Oligomeric states of putative active site mutants of HaCsx15. Related to Figures 2, 3 and 4.**

(A) SEC-MALS data for apo- tetramers H8A & R118A HaCsx15.

(B) Different active site conformations in HaCsx15-c$A_4$ tetrameric and filamentous forms.

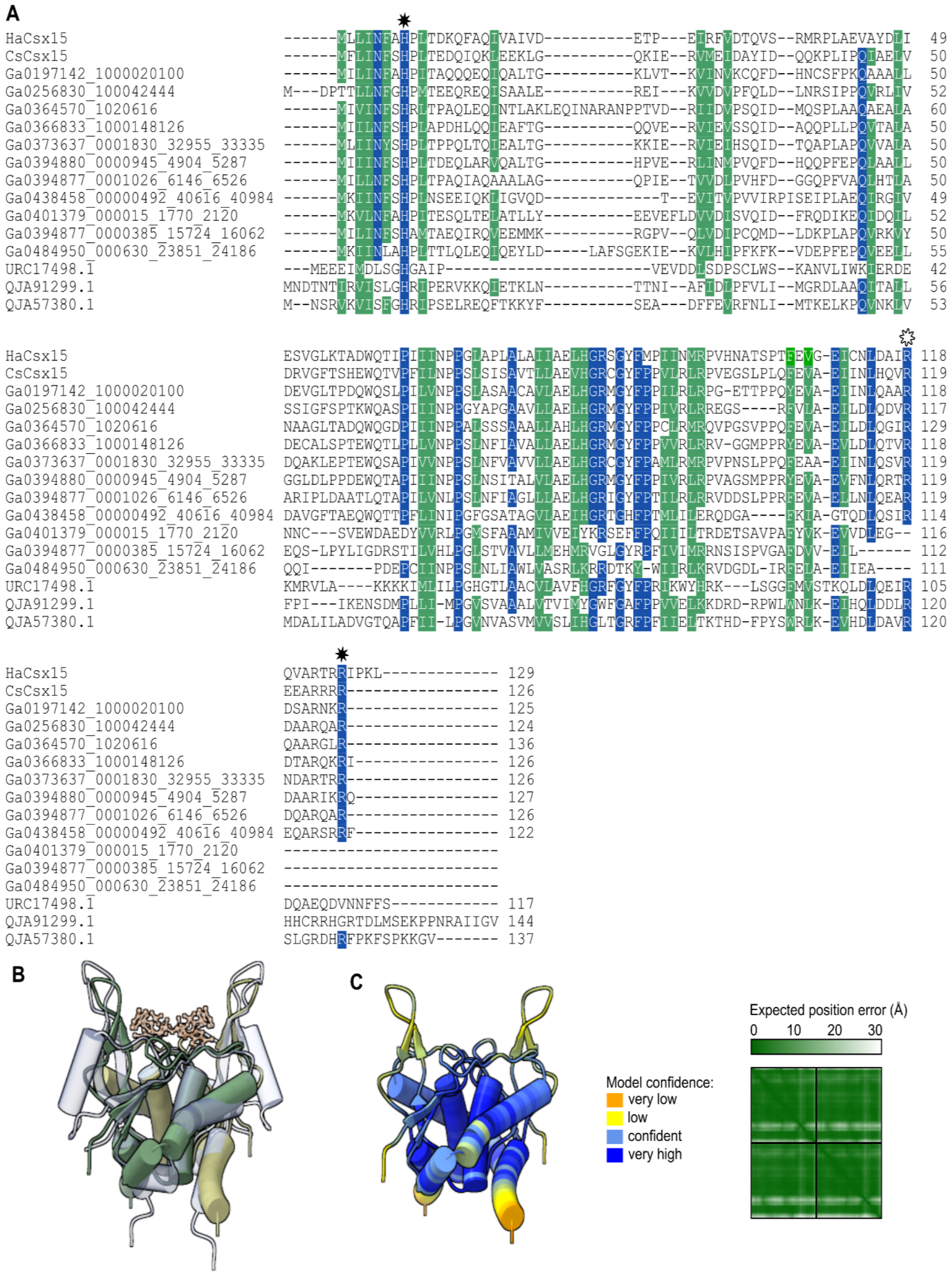


**Figure S8**. **Multiple sequence alignment of putative viral Csx15 homologs. Related to Figure 7.** (A) HaCsx15 and CsCsx15 are included for reference. Black stars above the alignment indicate active site residues in HaCsx15. White star indicates HaCsx15 R118 important for $cA_4$ binding. (B) The AlphaFold3 structural model of vCsx15 (NCBI: URC17498) (green), derived from the MAG genome of *Lokiarchaeota* virus Skoll Meg22_1214[21], is compared with the HaCsx15-$F_4cA_4$ structure (light violet). (C) The same model colored by the confidence (pLDDT) values is shown.

```
                <---------------------------------- Schlafen --------------------->
                                  *    * *
WP_015956247.1  MSRSLWTEIERWVINGTEGTKVEMKREFPLTDKPSRARLAKLIVAMANASGGTGYFIVGVVDKKD--RI-   67
MBL8150491.1    MSNSIWQEIQTWVINGVEGRKVEIKQEFALDDRQNRARFAKTITAIANAPVGKGYFIVGVVDIKS--RK-   67
MCD6167032.1    M--ELWDKLKRWILTGNESRKVELKQELPLKDRPSRAKLAKVVTSIANTPGGEGYIIVGVLEERH--RSD   66
MBN1815271.1    M-SRLWEILEHCIESGQEARKVDLKQLVDLASPDDRAKFAKDVAAMANTSGGPGYLVIGVVDRKH--RR-   66
MCB0259206.1    M--RLWEILKYYIETGQEARKVDLKQTLDLSSKKKRTEFAKDIAAMANTGGGSGYLIIGVLDQKH--REG   66
MBI5292449.1    M-DKLWRLLGNYVDSGRETRKVDLKREIEM--KAGAGKFAKDVAAIANTIGGTGYLIIGVIDLKE--RK-   64
MBI4771447.1    M-DRLWQILGPFVDGGRETRKVDLKYRMDLSSKGAQAEFAKDVSAIANTPGGAGYLVIGVQDAKA--RS-   66
WP_012302896.1  MNEAIWRRIKRYVDLGREAPKVELKQTLDLSTREKQAEFVKDVTAIANTPGGTGYVIIGVLDNTKCPRD-   69
HIE12021.1      MNVTIWRRIRRYVDLGHEAPKVDLKVTLDLSTKPNRAEFVKDVTAIANTPGGTGYLILGVWDNRKRPRVT   70
WP_170099038.1  MSDAIWRRIRRYVEQGHETPKVDLKLTVDLSKRDGRAEFAKDVTAIANTPGGTGYLIIGVLDSAACPRD-   69
HaCsx15         ----------------------------------------------------------------------

                ----------------------------------------------------------------------
WP_015956247.1  TNSIEEIIYGV-NDNLDSYQRLIQQALNEFTNPVPIVSYQEIIPPEISKKIGVIIIDRSRNKPHEITREC  136
MBL8150491.1    GTLLSEIVTGV-SYQPDDYQRAIQQALAEFTNPIPVVNYHHVSVPDVDKKIGVLIVERSRNRPHEITKES  136
MCD6167032.1    KNNLEEIIVGV-PYEHDAYQRLIQQALNEFTNPVPQVSYTEVLVPEVSRKIGVIVVERSRNQPHEIIQES  135
MBN1815271.1    SDDPSEYVVGFRPDDLEVFEQRMVQALENNCRPVPEIRYEEIRHPTVDRCIGVVMIPRGFSRPYTMNG--  134
MCB0259206.1    KDLLDDYVVGYTPDKLEVFEQQIVQALENNCSPVPEVHYEEINHPNIHHKIGVIVIPRSFSRPYIVNG--  134
MBI5292449.1    SNSPDDYVTGWNVTDLDGFSRWVNQSLSNYCNPIPRIEIEEMTHPVCGKKIGVIIIPRSLDCPHEIIRDG  134
MBI4771447.1    STAPEDYVPGWNVTDRDGFERGMNQALANYCNPPPGVELGEMVHPACGRRIGVIIIPRSFDRPHEVIREG  136
WP_012302896.1  RFLPENYVVGFSPSDPDELERQMIVALSNYCDPAPEIAFTQVLHPQTGRNIGVVTIARSAGRPHKFIRGS  139
HIE12021.1      YKKPEDYVPGFAPEDPDAFEREIIDALSHYCNRVPEVRFSQVMHPQTQRLIGVVIISRSSDKPHELIRSG  140
WP_170099038.1  HSTPEDYVVGFTHPNADNFYIQMIEALNQFCKPVPDIEYHEVVHEPTRRNVGVVTIKHSNMQPHSFIRPG  139
HaCsx15         ----------------------------------------------------------------------

                ------------------------->    <------------- Csx15 ------------------
                                                           *
WP_015956247.1  EKIKP-GIYLKRGSDVFQANREDILFM---TGANGKEQGIILNFT-HPVTENQIRQIEEKTGLHISEIVQ  201
MBL8150491.1    GDVKP-GIYIKRGAETFPAKREDILEM---TG-RSSQHAIIVNFT-HPITDEQIKQIEERERLYISEVVQ  200
MCD6167032.1    GDVRS-GIYVRRGAETFQARREELGFM---YG-SMSNYTVVVNFT-HPLSPHQIQQIEQQADVFIADVLA  199
MBN1815271.1    ------QIFIRRGSHTYPAAPDEVKQV---------PRRILVNFA-RAMDSTQIKQLEELLGAEIDEIID  188
MCB0259206.1    ------EIYIRRGTHTFRTSEDEVKQV---------NPRILINFA-REVDELQIEQLRKLLQADIDEIIN  188
MBI5292449.1    DGITK-GMYVRRGADTFTASREELRAM---MG--GTDVRIVLNFL-QPLTQLNREQIGQYLNARVYEVLE  197
MBI4771447.1    EGLRR-GIYTRRGADTFTASAAELREM---MG--AATALMVLNFH-QPFTLPQREQLERVTQSRVIEIIE  199
WP_012302896.1  TGIERQDVFIRRGTATFKASPQEIIEM---AQAPAFPEVIAINFSSHPLTSFQQEQLRERFHVHIEELIE  206
HIE12021.1      ERIEAPVVFVRRGTATYKATAEEIASMDREREVGPLSEVIVVNLSGHPLTDSQREELGAKYRVYVEELIE  210
WP_170099038.1  EGVQQHDVWIRRGTATFKATPEEIIKM---ATVGTSIPAVLINLSGHPLTEFQKTQLFDKYHVVVEELIE  206
HaCsx15         --------------------------------------MLLINFA-HPLTDKQFAQIVAI----VDETPE   27

                ----------------------------------------------------------------------
WP_015956247.1  PPKVPIHFNENNSFEDQVAKSVDEIALTDEQWQELNILVNVPGFATITAPLIAELHGRMGHFPKILRFKR  271
MBL8150491.1    PPKVPIHFLDDRSFEEQIRLEVDKIGLTEEEWQELPILINVPGFAYISAALIAELHGRMGHFPKLIRLRR  270
MCD6167032.1    PSKIPVHFDEVIPFEEQVKEIVDDVPLTDEEWQELSIIVNVPGYAVITAPLIAELHGRMGHFPKILRLRR  269
MBN1815271.1    ---VPGQLQDDQPYLPQARAMISAAGLTPAEWQSLPLVVNIHPFAPGAAAVLAWLHGLRGHFPDVVRMMR  255
MCB0259206.1    ---VPSKLFDNQPYLPQVRAMIDAVVLTPPEWQSLPLVVNIHPFAPAAAAVLAWIHGLRGFFPEIVRMAR  255
MBI5292449.1    PRSLPIRLDENLSFVEQAKILLNETGLTPEEWRTLPLIVNLPGFSPLSSILLAEMQGRMSRLPHIIRMRP  267
MBI4771447.1    PAGGPLRLDESRSFVLQARELLDAAGLTPEEWRTLPLVVNLPGFAPLAALLVAQLQGRMSRLPHVLRMRP  269
WP_012302896.1  ---LPVHFLAQEDLKPQAKKWIEEVGLALEEWSEKNIVLILPGLAPGAATILAAIHGLRGGFPKILWVYQ  273
HIE12021.1      ---LPVHFDPQQDLKSQAREYVHRIGLAPEEWSERQVHVILPGLAPGAAAILATIHGLSGGFPKVLWIYQ  277
WP_170099038.1  ---VPVHFDPTQELSPQIAKVIDDIDLAVEEWSERTIYVILPGLAPGAAAILAAIHGLKGGFPSILWIYQ  273
HaCsx15         IRFVDTQVSRMRPLAEVAYDLIESVGLKTADWQTIPIIINPPGLAPLALAIIAELHGRSGYFMPIINMRP   97

                ----------------------------------->
                                     ☼    *
WP_015956247.1  SSEDTSR-YDFEEVVQLQRIRDKARSRHQQ------  300
MBL8150491.1    LASDANR-YELAEIMQLQRIRDYARSRKQR------  299
MCD6167032.1    NADDPNH-YDFAEIIHLQRIRDRARERR--------  296
MBN1815271.1    NEQT-GH-FEVVEVLKLQSVRNEIRDWTVQL-----  284
MCB0259206.1    NEQN-NR-FEVVEVLLIQSVRNETRNLTVKFNK---  286
MBI5292449.1    TPADRTI-FEVAEVIALQDIRDKALEWSSGI-----  297
MBI4771447.1    SGADRTI-FEVAEVIELQRARDEAQQWVAGV-----  299
WP_012302896.1  APNDRTQ-YEVAQVVDLQALRDMGREIRSSVL----  304
HIE12021.1      APHDRTR-YELAQVVDLQECRDLAREVRVSVLGEPL  312
WP_170099038.1  SSQVGSQSYEIAQIIDGQSIRDTWRQTRTQTS---S  306
HaCsx15         VHNATSPTFEVGEICNLDAIRQVARTRRIPKL----  129
```

**Figure S9**. **Multiple sequence alignment of the Schlafen-Csx15 fusion proteins. Related to Figure 1.** The active site in HaCsx15 and the Schlafen active site motif (Exxxx(E/D)xK) in the fused proteins are indicated with black stars above the alignment. White star indicates HaCsx15 R118, important for $cA_4$ binding.

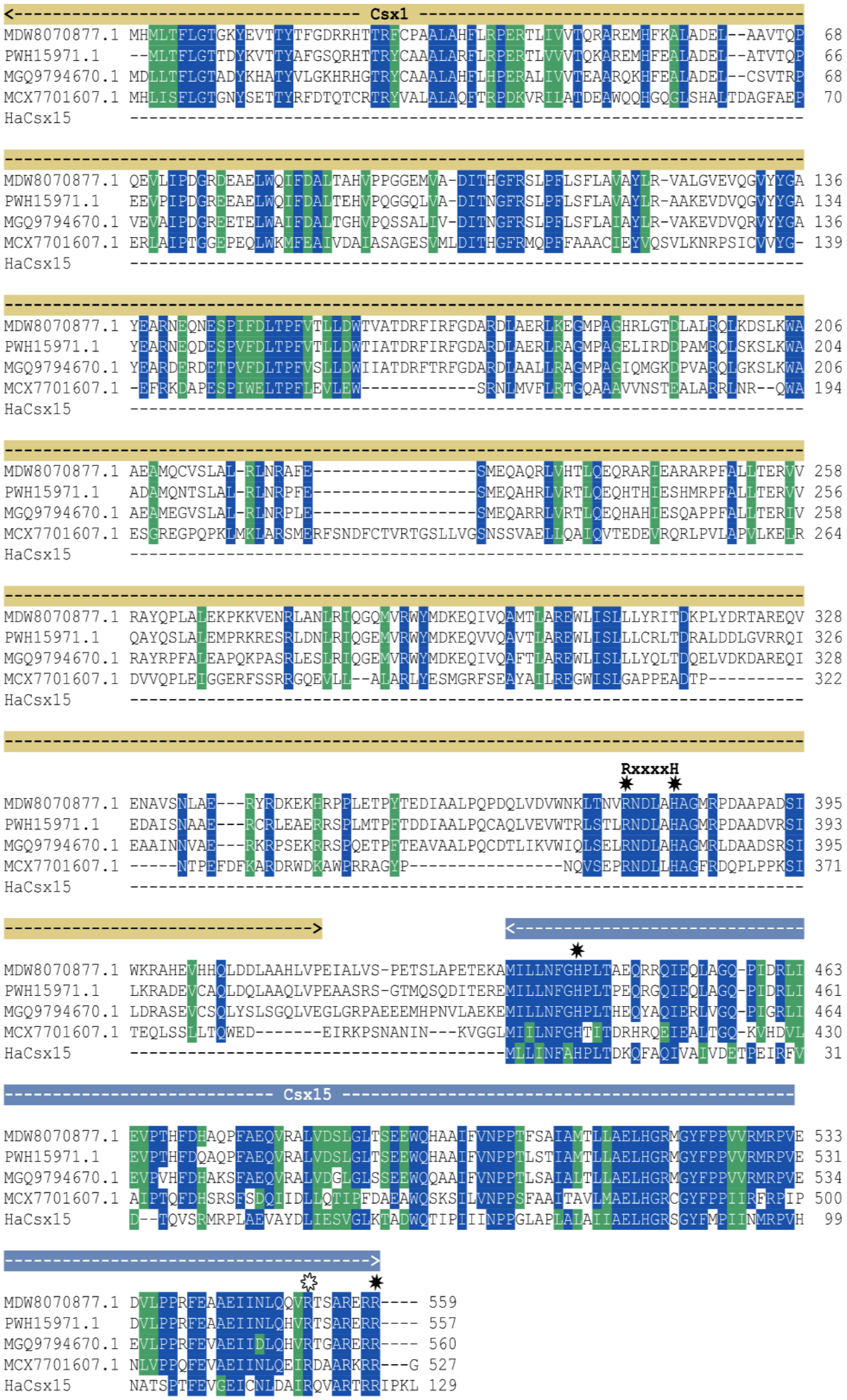


**Figure S10. Multiple sequence alignment of the Csx15 fusion with Csx1. Related to Figure 1.** Identical residues are shaded in blue, similar ones – in green. The HaCsx15 sequence is included for reference. Active site residues identified in HaCsx15 and the HEPN active site motif (RxxxH) in the fused Csx1 are indicated with black star above the alignment. White star indicates HaCsx15 R118, important for $cA_4$ binding.

**Table S4. Plasmids used in this study. Related to Figures 1, 2, 4 and 5.**

| Plasmid | Description | Usage |
|---|---|---|
| pBAD-HaCsx15 | pBAD-N-HSH-TEV-HaCsx15 ($Ap^R$) | Plasmid used for recloning the *Herpetosiphon aurantiacus* WT Csx15 variant into pMBP plasmid construct. |
| pBAD-CsCsx15 | pBAD-N-HSH-TEV-CsCsx15 ($Ap^R$) | Plasmid used for recloning the *Caldanaerobacter subterraneus* WT Csx15 variant into pMBP plasmid construct. |
| pMBP | pET-$His_{10}$-MBP-TEV ($Ap^R$) | Pasmid, used for cloning *Herpetosiphon aurantiacus* and *Caldanaerobacter subterraneus* WT Csx15 variants. |
| pMBP-HaCsx15 | pET-$His_{10}$-MBP-TEV-8aa-HaCsx15 ($Ap^R$) | Plasmid for expression of *Herpetosiphon aurantiacus* WT Csx15 variant fused with a TEV-cleavable $His_{10}$-MBP tag at the N-terminus. |
| pMBP-HaCsx15-H8A | pET-$His_{10}$-MBP-TEV-8aa-HaCsx15-H8A ($Ap^R$) | Plasmid for expression of *Herpetosiphon aurantiacus* H8A mutant Csx15 variant fused with a TEV-cleavable $His_{10}$-MBP tag at the N-terminus. |
| pMBP-HaCsx15-R118A | pET-$His_{10}$-MBP-TEV-8aa-HaCsx15-R118A ($Ap^R$) | Plasmid for expression of *Herpetosiphon aurantiacus* R118A mutant Csx15 variant fused with a TEV-cleavable $His_{10}$-MBP tag at the N-terminus. |
| pMBP-HaCsx15-R125A | pET-$His_{10}$-MBP-TEV-8aa-HaCsx15-R125A ($Ap^R$) | Plasmid for expression of *Herpetosiphon aurantiacus* R125A mutant Csx15 variant fused with a TEV-cleavable $His_{10}$-MBP tag at the N-terminus. |
| pMBP-CsCsx15 | pET-$His_{10}$-MBP-TEV-8aa-CsCsx15 ($Ap^R$) | Plasmid for expression of *Caldanaerobacter subterraneus* WT Csx15 variant fused with a TEV-cleavable $His_{10}$-MBP tag at the N-terminus. |
| pMBP-CsCsx15-H8A | pET-$His_{10}$-MBP-TEV-8aa-CsCsx15-H8A ($Ap^R$) | Plasmid for expression of *Caldanaerobacter subterraneus* H8A mutant Csx15 variant fused with a TEV-cleavable $His_{10}$-MBP tag at the N-terminus. |
| pBAD_AvCami1_S11A | pBAD_AvCami1_S11A ($Ap^R$) | Plasmid, used for recloning the *Allochromatium vinosum* ring nuclease-dead Cami1 variant into pETDuet-1 plasmid construct. |
| pAvCami1_S11A | pETDuet-1_AvCami1_S11A ($Ap^R$) | Plasmid, expressing *Allochromatium vinosum* ring nuclease-dead Cami1 variant. Used in the *E. coli* survivability assay. |
| pAvCami1_S11A_HaCsx15 | pETDuet-1_AvCami1_S11A_HaCsx15 ($Ap^R$) | Plasmid, expressing *Allochromatium vinosum* ring nuclease-dead Cami1 and *Herpetosiphon aurantiacus* WT Csx15 |

| | | |
|---|---|---|
| | | variants. Used in the *E. coli* survivability assay. |
| pAvCami1_S11A_HaCsx15-H8A | pETDuet-1_AvCami1_S11A_HaCsx15-H8A ($Ap^R$) | Plasmid, expressing *Allochromatium vinosum* ring nuclease-dead Cami1 variant and *Herpetosiphon aurantiacus* WT Csx15 variant. Used in the *E. coli* survivability assay. |
| pCsm | pCas/Csm_dCsm3_dHDCas10_ΔCsm6′ΔCsm6 ($Str^R$) | Plasmid for expression of protein subunits of *Streptococcus thermophilus* nuclease-dead Csm complex. Used in the *E. coli* survivability assay. |
| pCRISPR | pACYCDuet-1_Leader-[R-mut1GFP]4-TR ($Cm^R$) | Plasmid for expression *of* crRNA complementary to *Aequorea victoria* GFP mRNA. Used in the *E. coli* survivability assay. |
| pTarget | pCOLADuet-1_GFP ($Kn^R$) | Plasmid for expression of *Aequorea victoria* GFP gene. Used in the *E. coli* survivability assay. |
| pEmpty | pETDuet-1 ($Ap^R$) | Control plasmid used in the *E. coli* survivability assay. |

* $Str^R$ – streptomycin-resistance, $Ap^R$ – ampicillin-resistance, $Cmr^R$ – chloramphenicol-resistance, $Kn^R$ – kanamycin-resistance.
